# Upper bill bending as an adaptation for nectar feeding in hummingbirds

**DOI:** 10.1101/2024.10.01.615288

**Authors:** Alejandro Rico-Guevara, Diego Sustaita, Kristiina J. Hurme, Jenny E. Hanna, Sunghwan Jung, Daniel J. Field

## Abstract

Observations of maxillary (upper bill) bending in hummingbirds have been considered an optical illusion, yet a recent description of out-of-phase opening and closing between their bill base and tip suggests a genuine capacity for bill bending. We investigate bill kinematics during nectar feeding in six species of hummingbirds. We employed geometric morphometrics to identify bending zones and combined these data with measurements of bill flexural rigidity from microCT scans to better understand the flexing mechanism. We found that the mandible remains in place throughout the licking cycle, while the maxilla undergoes significant shape deformation, such that the distal portion of the upper bill bends upwards. We propose that bill bending is a key component of the drinking mechanism in hummingbirds, allowing the coordination of bill function (distal wringing and basal expansion) and tongue function (raking/squeegeeing) during intraoral transport. We present a fluid analysis that reveals a combination of pressure-driven (Poiseuille) and boundary-driven (Couette) flows, which have previously been thought to represent alternative drinking mechanisms. Bill bending allows for separation of the bill tips while maintaining a tightly closed middle section of the bill, enabling nectar exploitation in long and narrow flowers that can exclude less efficient pollinators.

## 1. Introduction

Despite often exhibiting lightweight, pneumatized skeletons, the bones of birds are usually extremely rigid [1]. Bone flexibility is generally limited in most tetrapods [2]; however, some groups of birds exhibit flexible bones in their skulls for functional reasons (e.g., [3–5]). The bill is one of the most evolutionarily labile external structures in birds, reflecting the immense disparity of feeding styles exhibited across roughly 11,000 extant species. In addition to this remarkable ecomorphological disparity, the striking dexterity of the avian bill is enhanced by variable degrees of cranial kinesis, the origin of which may have helped compensate for the loss of manual dexterity associated the origin of avian flight [6], and at least an incipient degree of cranial kinesis appears to have been present in the last common ancestor of all extant birds [7]. The most common form of avian cranial kinesis involves movement of the entire upper bill with respect to the cranium by flexion of a region of narrow bone where the beak meets the cranium, known as the cranio-facial or nasofrontal hinge [8–13]. However, some bird clades exhibit additional specialisations for bill flexion in which the very bones of the bill are capable of bending, such as lateral bowing of the lower jaws (e.g., [4,14,15]) and dorsiflexion of the upper jaws (i.e., rhynchokinesis; [16,17]).

Although all birds possess skulls that are kinetic to an extent [16], rhynchokinesis near the bill tip is especially well-developed in shorebirds (Charadriiformes: Scolopacidae). This enables many shorebird species to open and close the tips of the bill independently of the bill base, which is generally associated with their substrate-probing feeding habits [16,18,19] but has also been shown to help facilitate the unique surface-tension transport mechanism of phalaropes [20,21], and enhance feeding efficiency on small aquatic prey in Calidris sandpipers [22,23]. However, bill bending is considerably less well understood outside of shorebirds, and few other avian clades are known to exhibit this specialisation. An additional group capable of bill bending is Strisores, which in part includes nightjars (Caprimulgidae) and Apodiformes (e.g., swifts and hummingbirds) [24–26]. Most subclades within Strisores are aerial insectivores that employ mandibular bending for gape expansion [14], and the lower jaws of hummingbirds have been observed to bend ventrally and laterally during flycatching manoeuvres and visual displays, indicating considerable flexibility of their lower jaws [15,27,28]. Hummingbird ancestors were short-billed aerial predators resembling modern cypselomorphs such as swifts (Apodidae; [29,30]. Through coevolution with increasingly specialised ornithophilous plants, hummingbird beaks became long and slender [30]. To be able to take advantage of nectar as a food resource without damaging the flower (allowing continued nectar production), hummingbird tongues became elongate and protrusible, evolving as highly efficient liquid-collecting devices closely coupled with, and paralleling, elongation and thinning of their bills to fit inside narrow flowers (reviewed by Rico-Guevara et al. [31]). Interestingly, such a high degree of morpho-functional specialisation for nectarivory has not compromised their ability to effectively perform aerial insectivory (the plesiomorphic dietary state for Strisores), in large part because of mandibular bending. Yanega and Rubega [15] demonstrated that ventral flexion of the mandible (lower jaw) enhances aerial prey capture success. Here, we evaluate for the first time whether another form of bill bending—distal rhynchokinesis of the upper jaw—exists in hummingbirds and whether it confers advantages for enhancing drinking efficiency.

From a mechanical perspective, “drinking” is simply an animal’s behaviour of transporting fluid into the body. There are two fundamental modes of fluid transport: pressure-driven (Poiseuille) and boundary-driven (Couette) flows [32–34]. Most animals employ a pressure-driven mechanism (suction) using a confined oral structure, while a few species (mostly carnivoran mammals) employ boundary-driven flow (lapping) [35–37]. For small animals, capillary-driven flows can be used, which can be categorised as pressure-driven flow to some extent [38–41]. Although widely hypothesised [42], capillary filling of the tongue during nectar feeding in birds has only been documented in Pied honeyeaters [43], while other honeyeater species (Passeriformes: Meliphagidae) employ a similar mechanism to hummingbirds, the most intensively studied group of avian nectarivores [recent reviews in [42–44]].

Previous research on hummingbird drinking biomechanics has shown how the tongue collects nectar [45,46], and how this liquid is offloaded inside the bill [45]. Briefly, during tongue protrusion, hummingbirds flatten the distal grooves of their tongues [45,46], which are hollow to collect the nectar but that comprise only about half of their tongue length [47], so they cannot be used as straws. While the tongue moves through the air during protrusion, the elastic energy loaded into the groove walls during the flattening is conserved by a remaining layer of liquid inside the grooves acting as an adhesive [45]. When the tongue touches the nectar, the fringed tip opens up [48] and the supply of fluid allows the release of the elastic energy which expands the grooves and pulls the nectar to fill the entire tongue grooves [45]. The tongue is then retracted, the fringes at the tip capture fluid via a natural origami self-folding trap [48], and the tongue, now sealed at the tip and completely filled with nectar, is quickly brought inside the oral cavity while the bill tips are kept separated. As soon as the tongue is inside the bill, the bill tips close leaving only a small gap through which the tongue grooves are extruded through, wringing the nectar inside the oral cavity near the bill tip, flattening the tongue, and starting the process again with the next cycle [45], with a frequency of around 15Hz [31]. In spite of these advances in our understanding of the nectar collection mechanisms, however, the process by which nectar is transported to the throat during hummingbird feeding has only recently been described [49], and the underlying mechanics for the bill motions that foster this intraoral transport are unknown. The default expectation for bill separation, set by a lever mechanism wherein the jaws hinge upon the naso-frontal and quadrato-mandibular joints at the bill base [5], is that separation at the base is followed by in-phase synchronised (same period) separation of the tip—the same way in which a pair of scissors works. Hummingbirds, on the contrary, exhibit phase-shifted (out of phase) separation between the base and the tip, with the bill’s middle portion showing the least amount of separation [49], akin to the motion of a seesaw. However, the mechanism by which this phase-shifted bill opening operates has yet to be explained. It could potentially be achieved by different mechanisms such as mandibular bending (lower bill tip deflecting downwards), mandibular rotation (a curved lower jaw elevated basally and hinging in the middle of the upper jaw), maxillary bending (upper bill tip deflecting upwards), or combinations of the above mechanisms. Nitzsch [50] first observed hinges in the still flexible bills of juvenile hummingbirds and inferred that hummingbirds could use bending of the tip of the maxilla to wring nectar off the tongue during extrusion. Such bill bending might optimise the feeding process through coupling of bill and tongue movements (see [51–53]), and although this was never experimentally demonstrated, Bühler [14] also suggested potential flexion zones along the maxilla.

Despite these early studies hypothesising a role for bill bending during hummingbird feeding, Zusi’s [16] exhaustive comparative survey of avian rhynchokinesis (i.e. bending of the upper bill) found no osteological evidence supporting distal bill bending in hummingbirds. Zusi [16] therefore concluded that most species of hummingbirds were proximally rhynchokinetic, and a few actually prokinetic, meaning that any dorsally directed bending of the upper jaw would be restricted to the base of the bill. However, Zusi [17] later acknowledged the slight bending of the tip of the upper jaw of hummingbirds observed in videos by Rico-Guevara and Rubega [48] and concluded that distal rhynchokinesis was not only possible but could even facilitate nectar consumption [17]. Furthermore, Zusi [17] underscored the need for “precise measurements of cranial kinesis of live hummingbirds and knowledge of the physical properties of the structural components of the prepalatal upper jaw.”

Here, we tested the hummingbird bill-bending hypothesis by combining high-speed videography, geometric morphometrics, and micro-computed tomography to quantify bill bending in- and ex-vivo across a wide phylogenetic sample of hummingbirds (Strisores: Trochilidae). Our objective was to quantify upper bill flexibility of hummingbirds during nectar feeding, and to determine whether the hummingbird upper jaws exhibit a distinct dorsoventral flexural rigidity profile that facilitates bending of the distal portion of the bill. These insights help clarify the specialised form of nectar drinking exhibited by the ∼350 species of extant hummingbirds.

## 2. Methods

### 2.1. Video analyses

We filmed free-living hummingbirds trained to drink nectar from artificial feeders with clear sides, which were wide enough to avoid limiting any bill tip motion during feeding (e.g., [46,48]), in Colombia, Ecuador, and the United States (all at private locations with landowner permission). We worked with adult birds (judging by the absence of many visible corrugations at the bill base [54–56]) drinking while hovering (e.g., Movie S1), with high-speed cameras (TroubleShooter HR, Fastec Imaging, 1000 frames/s, 1280 × 512 pixels) coupled with macro lenses (Nikon 105mm f/2.8 VR). All filming activities were reviewed and authorised by the Institutional Animal Care and Use Committee at the University of Washington (Protocols 4498-01, 4498-03) and at the University of Connecticut (Exemption Number E09-010). We processed videos from representative subjects of six species of hummingbirds belonging to four of the nine major hummingbird subclades to maximise phylogenetic, morphological, and functional diversity in our sample for assessing the phylogenetic distribution of bill bending in Trochilidae (Table S10). For each of the six subjects, high-speed videos of licking footage (e.g., Movie S1) were converted into a series of image files using QuickTime Player Version 7.6.6. All images in each sequence were digitally and equally enhanced by consistently adjusting the contrast and brightness using ImageJ 1.45p [57], to maximise the visibility of targeted landmarks (dorsal culmen curvature and tomium).

We measured ten licking cycles (i.e., “licks”) per individual (e.g., Movie S1), with the exception of *Chalybura buffonii*, for which only five cycles were analysed. We defined the start of a lick as the first frame in which the tongue is visible while being protracted, and the end of each lick as the last frame before the tongue is visible again during protraction. The duration of one lick was usually less than 1/10th of a second, and we extracted a subset of eleven equally spaced frames from each high-speed video sequence to analyse (yielding ten time-steps about ten milliseconds apart). For the analyses, the first (“frame 0”) and eleventh (“frame 10”) frames were the ones directly preceding the tongue’s first appearance during protraction, thus encompassing a full cycle.

### 2.2. Assessment of bill deformation

Starting with the lateral views and for each of the eleven frames per lick, we digitally traced three bill contours to create profile lines using tpsDIG2 [58]. The first line followed the culmen beginning distally at the maxillary tip and ending at the most proximal point of the exposed culmen (the distal-most extent of feathering). A vertical line perpendicular to the bill axis was followed ventrally from the most proximal point of the exposed culmen, and its intercepts with the other bill contours were used to place points defining the most proximal points of the other profile lines: one on the maxillary tomium, and one on the mandibular ventrum (figure 1A). We then traced the maxillary tomium profile from this proximal point (intercept with the end of exposed culmen perpendicular line) to the maxillary tip, as well as the profile of the mandibular ventrum (defined here as the lower jaw contour, that is, the exposed culmen’s counterpart on the lower jaw) from the most proximal point as defined above to the mandibular tip. These three bill profile lines were then resampled using tpsDIG2 so that each line had 21 equidistant semilandmarks, with which we could assess bill deformation (figure 1B). Landmark tracking was ultimately performed by a few different observers, so we conducted a repeatability experiment (electronic supplementary materials S1) to assess the potential impacts of within- and among-observer tracking error, in a manner similar to that described by Moccetti et al. [59].

**Figure 1.**
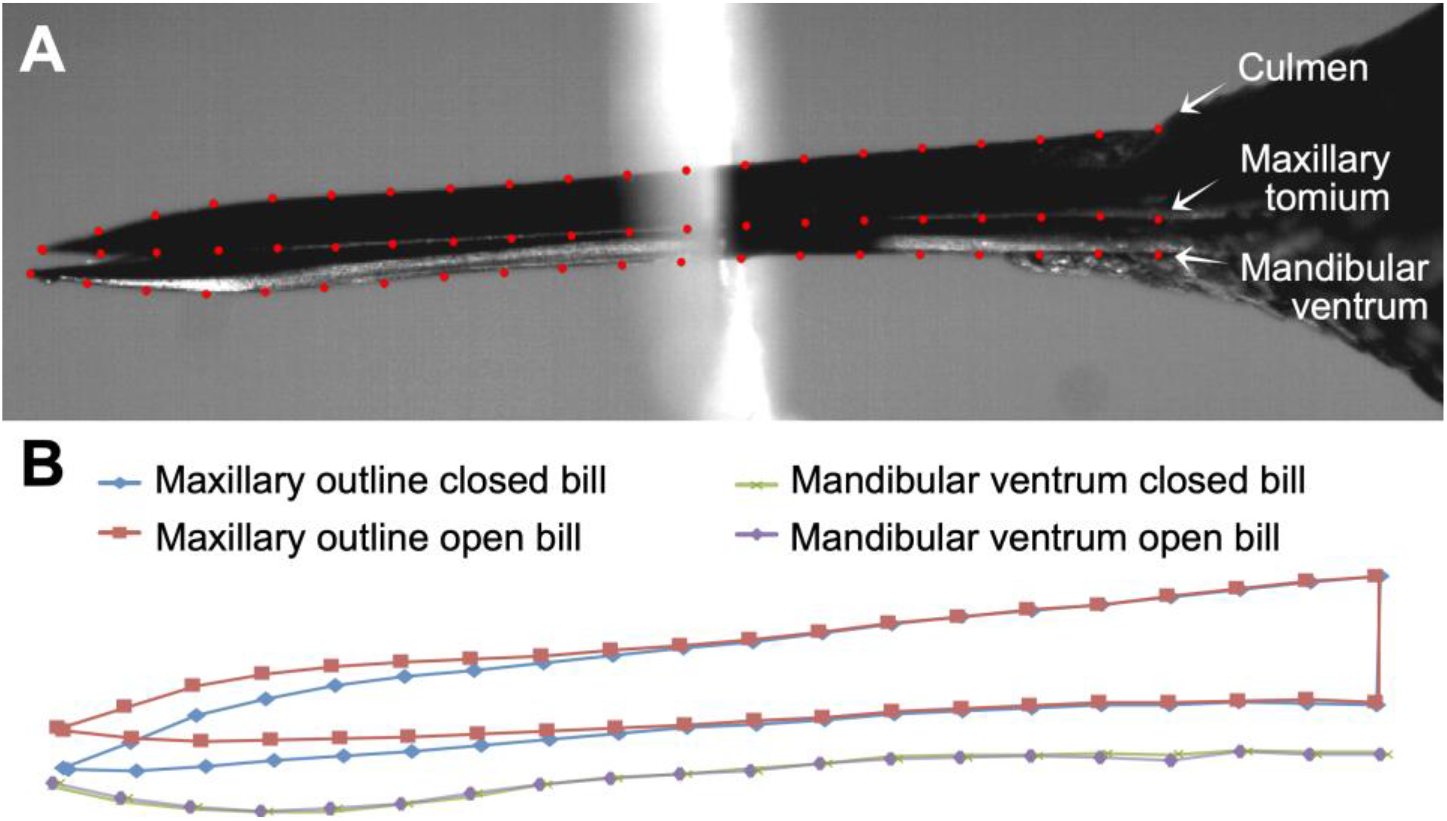
A) Description of semilandmarks used for video analyses. B) Outputs from a geometric morphometric analysis of the landmarks showing the most extreme displacements of the upper bill (blue and red) and the visible bottom of the mandible (green and purple) during a given licking cycle.

We used a geometric morphometric approach to visualise the relative displacements of landmarks throughout a given licking cycle, using approaches adapted from [60–62]. This analysis allowed us to assess the deformation of the culmen (underlain by the premaxillary, nasal, maxillary, and jugal bones) during the course of a lick, to help identify the bill’s bending zones. We used tpsRelw32 [63] to perform generalised Procrustes alignment (GPA; using generalised least squares) of all the frame image landmark configurations (“frames,” from here on) of all lick sequences (collectively) for each subject filmed. We then performed a relative warps analysis (i.e., principal component analysis of the partial warp scores [63]) to validate the observed culmen deformations. For this analysis we chose to slide semilandmarks (by resampling along the contour lines) using the method that minimises the Procrustes distance (d2), both based on recommendations of [64], as well as our own observations that a minimum bending energy approach seemed to introduce artefacts. We used Procrustes ANOVA, a non-parametric approach that leverages residual randomization permutation procedures, as described by Adams and Collyer [65,66] and Collyer and Adams [67], implemented in function procD.lm of the R package geomorph [68]. This procedure tests for the effects of independent variables on the full set of dependent (shape) variables (i.e., the Procrustes coordinates: the landmark coordinates aligned by generalised Procrustes analyses). We performed Procrustes ANOVAs for each subject to test the significance of the changes in bill shape observed across sequential frames. For these analyses, the effects of “dataset” (in the intra- and inter-observer landmark repeatability analysis; electronic supplementary materials S1) and “cycle” (in the omnibus test of bill deformation for each species) were included as blocking factors. One advantage of this approach is its robustness to the unfavourably high p:n ratios inherent to geometric morphometric datasets [67]. Generalised least-squares analyses were performed using Type I Sums of Squares and Cross-products and were set to 9,999 permutations of null model residuals. Wireframe plots for selected sequential frames were generated using the plotRefToTarget function of R package geomorph [68], with magnification set to 1.5X-7X to better visualise the shape changes.

We summarised the total GPA variances (S^2^) for each landmark along the dorsal culmen and tomium. We felt that using the GPA variance (which provides a measure of the variability of the x and y positions of each Procrustes-aligned landmark) was appropriate and sufficient to fulfil our objectives of: (1) demonstrating upper bill movement at each landmark during a licking cycle, and (2) generating a quantitative variable for the relative displacements of individual landmarks along the length of the culmen over time to juxtapose with the flexural rigidity data described below. We acknowledge the potential shortcoming of this approach, in that during the Procrustes alignment procedure, the variance in landmark positions is distributed across all landmarks [69]. However, our preliminary investigation of this effect (by randomly varying pixel positions at some landmarks of a single frame within the ranges of x and y displacements observed during a lick cycle), indicated that the variance attributed to other (unvaried) landmarks is negligible, amounting to ≤ 5.7 % (depending on the number of landmarks varied) of the variance in altered landmark positions, and < 1% of the total landmark variances observed during an actual lick cycle. We chose not to slide the semilandmarks for this particular analysis, primarily because we wanted to observe the actual variation in landmark positions over the course of a lick cycle while avoiding any potential influence of statistical artefacts, as the sliding procedure tends to exaggerate variation in the vertical dimension (e.g. [64]), which is the primary locus of the signal that we endeavoured to detect.

### 2.3. High-resolution X-ray computed tomography (µCT)

We generated µCT scans (down to 5-micron voxel resolution) of hummingbird heads from specimens from the Yale Peabody Museum corresponding to the same species studied, to discern between bone and rhamphotheca and thus match our high-speed videos to the bone density analyses (see below). The specimens were scanned at The University of Texas High-Resolution X-ray Computed Tomography Facility, Austin, Texas, and on aµCT35 instrument (Scanco Medical AG) at the Yale Core Centre for Musculoskeletal Disorders, New Haven, Connecticut. Scans were performed at 70 kV and 10W, with Xradia 0.5 and 4X objectives, and 1 mm SiO2, or no filter. Specimens were scanned in three parts, with the scans stitched together using Xradia plugins, and voxel size was between 15.5 and 5.2 μm. 16bit TIFF images were reconstructed in Xradia Reconstructor, and the total number of slices per specimen was between 2,223 and 2,854, with scan times between 4 and 7 hours.

### 2.4. Dorsoventral flexural rigidity analysis

A longstanding problem in vertebrate functional morphology has been the difficulty associated with quantifying the flexural rigidity profile of bones across their entire length. Although subjecting bones to mechanical tests to determine bending resistance provides a quantitative estimate of flexural rigidity at certain points along a bone, such tests provide little information about a bone’s overall mechanical design. A method of non-destructive flexural rigidity determination, BendCT, facilitates the quantification of a bone’s flexural rigidity profile along a user-defined bending axis across its entire length [4]. We used BendCT to test the hypothesis that the hummingbird upper jaw’s “bending zone” is characterised by a marked decrease in the bone’s dorsoventral flexural rigidity, relative to proximal and distal rigid zones.

Flexural rigidity is the product of an object’s elastic stiffness, quantified as the Young’s modulus (*E*), and its cross-sectional shape, quantified as the second moment of area (*I*):

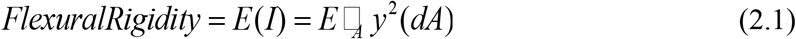

where *y* is the distance between the area increment *dA* and the bending axis of interest. In bone, mineral density is the main determinant of *E*, allowing *E* to be estimated using equations relating experimentally derived values of *E* to non-invasive proxies of bone mineral density (e.g., Hounsfield units). Calibration of a Hounsfield unit-mineral density relationship was accomplished by scanning hydroxyapatite phantoms of known mineral density under the same conditions as the hummingbird specimens. We then followed previous work (e.g., [2,4,70] by applying the regression derived by Ciarelli et al. [71] to estimate *E* averaged across all three orthogonal directions from mineral density. This approach ignores the potential influence of bone water content and material anisotropy on *E*. However, *EI*, using average values, does not take into account non-homogeneity in cross-sectional mineral density, which leads to regional variation in *E*. To address this issue, BendCT computes cross-sectional flexural rigidity pixel-by-pixel for each serial cross-section, by defining *dA* as the area of a pixel, and taking the summation form of the integral as follows:

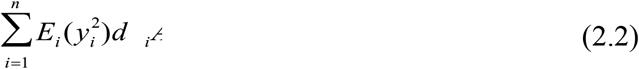

where *i* identifies an individual pixel, and *n* is the total number of pixels in a user defined region of interest. Thus, BendCT facilitates the quantification of flexural rigidity weighted by the effect of bone density on *E*, relative to the dorsoventral bending axis of the hummingbird upper jaw. Stacks of high-resolution microCT images were obtained for the upper jaws of the hummingbirds for which performance data were collected, from the nasofrontal hinge to the rostral tip of the bill. BendCT was used to reconstruct the pattern of dorsoventral flexural rigidity variation throughout the hummingbird upper jaw, to test the hypothesis that hummingbird upper jaw bending is facilitated by depressed dorsoventral flexural rigidity within the jaw’s bending zone. This hypothesis predicts that regions of relatively high dorsoventral flexural rigidity should flank the bending zone at the bill’s distal and proximal ends; however, the bending zone itself should be characterised by a pronounced reduction in dorsoventral flexural rigidity. To examine this, we paired the dorsoventral flexural rigidity data (based on the combined frontal process of the premaxillae, defined as ‘top’, and the main body of the premaxillae, defined as the ‘fused’ portions of the bill only —see [17] for a discussion on the anatomical nomenclature of the hummingbird bill bones—, to account for resistance to bending in the dorso-ventral axis) with the kinematic geometric morphometrics data. Because the subjects filmed and the specimens µCT scanned were not the same individuals, we aligned the BendCT flexural rigidity data with the kinematic data on their own sets of x and y axes to assess the congruence between datasets. We scaled the lengths of the bills of the specimens from which the dorsoventral flexural rigidity data were obtained to the total culmen lengths estimated for each subject (i.e. from rostral bill tip to nasofrontal hinge), based on their measured exposed culmen lengths (i.e., from the rostral bill tip to the cranial extent of the rhamphotheca) and the relationship between exposed culmen and total culmen lengths measured from other specimens of the same species. This alignment of the datasets takes into account that the BendCT data distally-to-proximally starts on the bony part of the bill, while the geometric morphometrics data starts at the keratinous rostral tip of the culmen. We then used the CT scans themselves to measure the portion of the bill tip that is only keratin, resulting on a more accurate alignment.

## 3. Results

### 3.1. Rhynchokinetic bill deformation in live hummingbird subjects

The GPA landmark variances during lick cycles exceeded frame error variances (based on non-overlapping 95% CIs; electronic supplementary materials, S1.1) along landmark positions 1–2, 8–12, and 14–18 in *Calypte anna* (figure 2), indicating zones of mobility, or ‘bending zones’, where landmark variances were greater than tracking error (e.g., landmark positions 1-2 and 8-12; figure 2) punctuated by regions of no discernible mobility where variances were actually lower than error (e.g., landmark positions 4-6 and 14-18; figure 2). Based on the (conservative) error analyses of this subject, we are confident that the regions of high and low GPA variances in other subjects accurately reflect the zones of greatest and least mobility (respectively) along the lengths of their bills.

**Figure 2.**
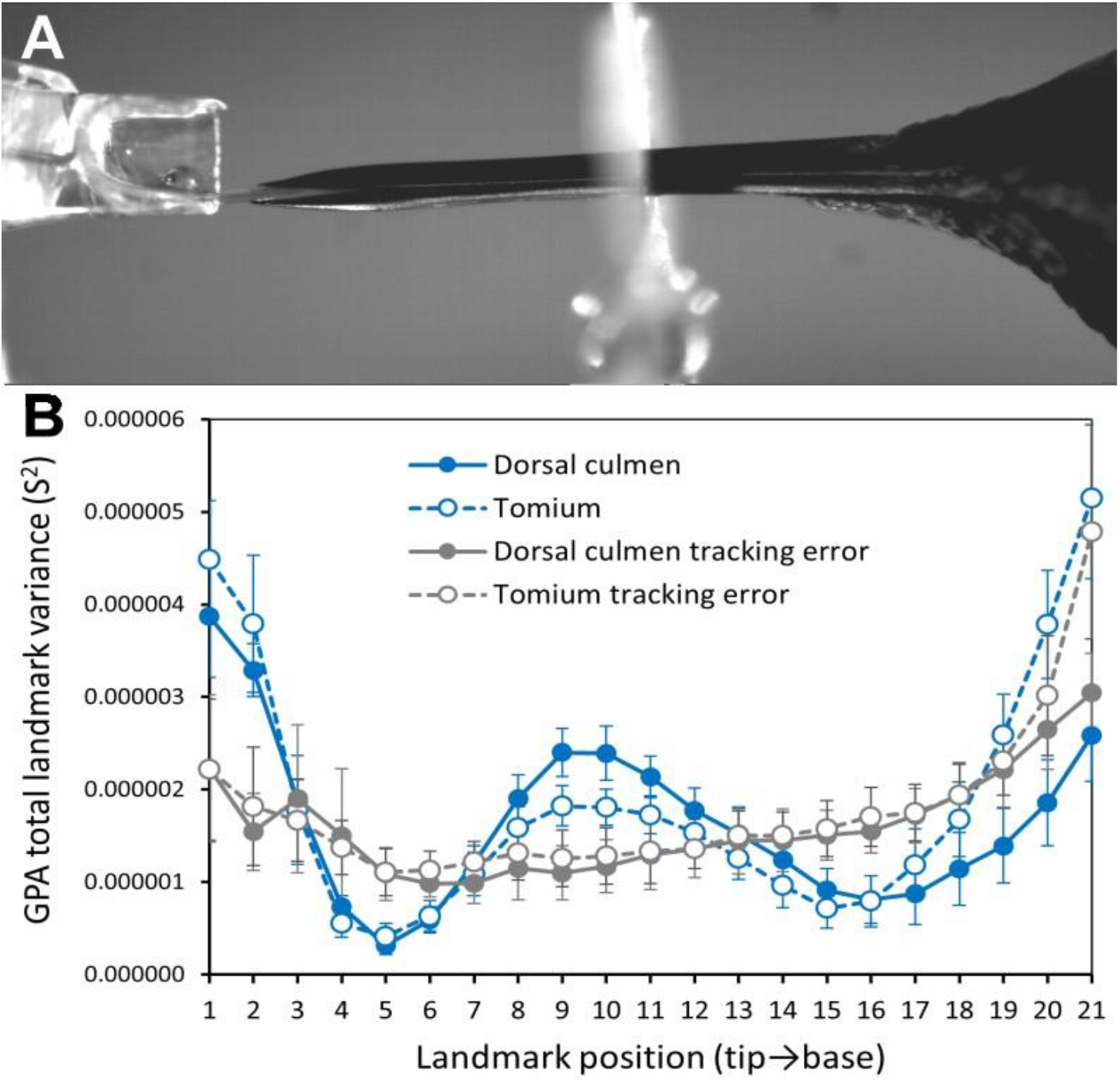
A) Image from a *Calypte anna* lick cycle. B) Mean ± 95% CI (*n* = 10 lick cycles) GPA total landmark variances (S^2^) computed over 11 sequential frames, for the dorsal culmen (blue filled symbols and solid line) and tomium (blue open symbols and dashed lines) of the *Calypte anna* subject. The mean ± 95% CI (*n* = 11 frames) GPA landmark variances due to tracking error obtained from a single lick cycle tracked by five different participants three times each (electronic supplementary material, S1.1) are shown in grey (dorsal culmen filled symbols and solid line; tomium open symbols and dashed line).

The shapes of the culmens changed during licking as evidenced by their trajectories through morphospace across sequential images (figures 3, 4; figure S3). The percentages of variance explained by the first two PC axes varied considerably among subjects/taxa, but in each subject the changes in bill shape followed similar trajectories across replicate lick cycles. The Procrustes ANOVAs for each subject revealed that, although the variation among lick cycles was significant, cycle accounted for a relatively small proportion of the variance (*R*^*2*^ = 0.08–0.25) compared to the significant shape changes detected across frames (*R*^*2*^ = 0.20–0.77; table S2–S7). However, the degree of deformation in the culmen varied considerably among subjects/taxa, largely owing to differences in overall bill shapes (table S8; figure S10). In general, the culmen exhibited the greatest deformation along the rostral third of the bill, secondarily along the middle third, and to varying extents at the base (figure S3). Principal component 1 consistently expressed some degree of relative dorsoventral narrowing vs. widening of the culmen, along with a relative straightening vs. a relative decurvature of the rostral half of the culmen (figure 3, 4), from one direction of the PC axis to the other (note, however, that the directions were determined arbitrarily for each separate analysis). Whereas PC1 (which explained the bulk of the variance of each data set) captured most of the “bill bending” deformation among sequential frames, changes along PC2 were more subtle, and typically reflected comparatively minor differences in shape among licking cycles (figures S4–S9).

**Figure 3.**
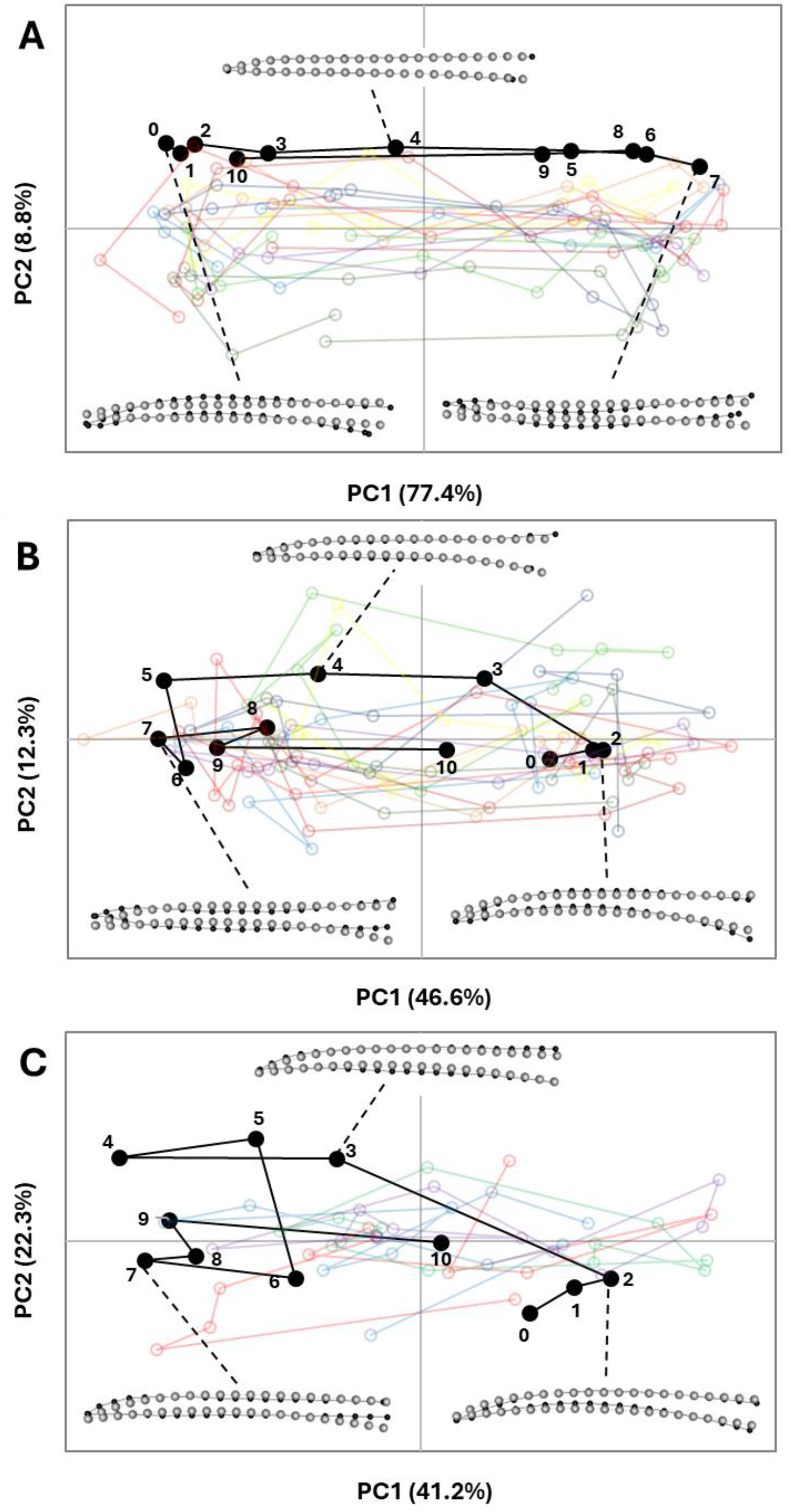
Shape changes in PC1-PC2 morphospace observed in the culmen of species with the most similar, relatively straight, overall bill shapes for A) *Calypte anna*, B) *Amazilis amazilia*, and C) *Chalybura buffonii*, during lick cycles. In each graph, “Lick01” (black filled symbols, numbered sequentially) is shown for reference; the semi-transparent open symbols and traces in other colours are those of the other nine lick cycles (Lick02-10). The insets show representative shape changes (black dots and lines; relative the grey dots and lines of the consensus configuration, magnified by 7X) for selected frames during Lick 01 for each species.

**Figure 4.**
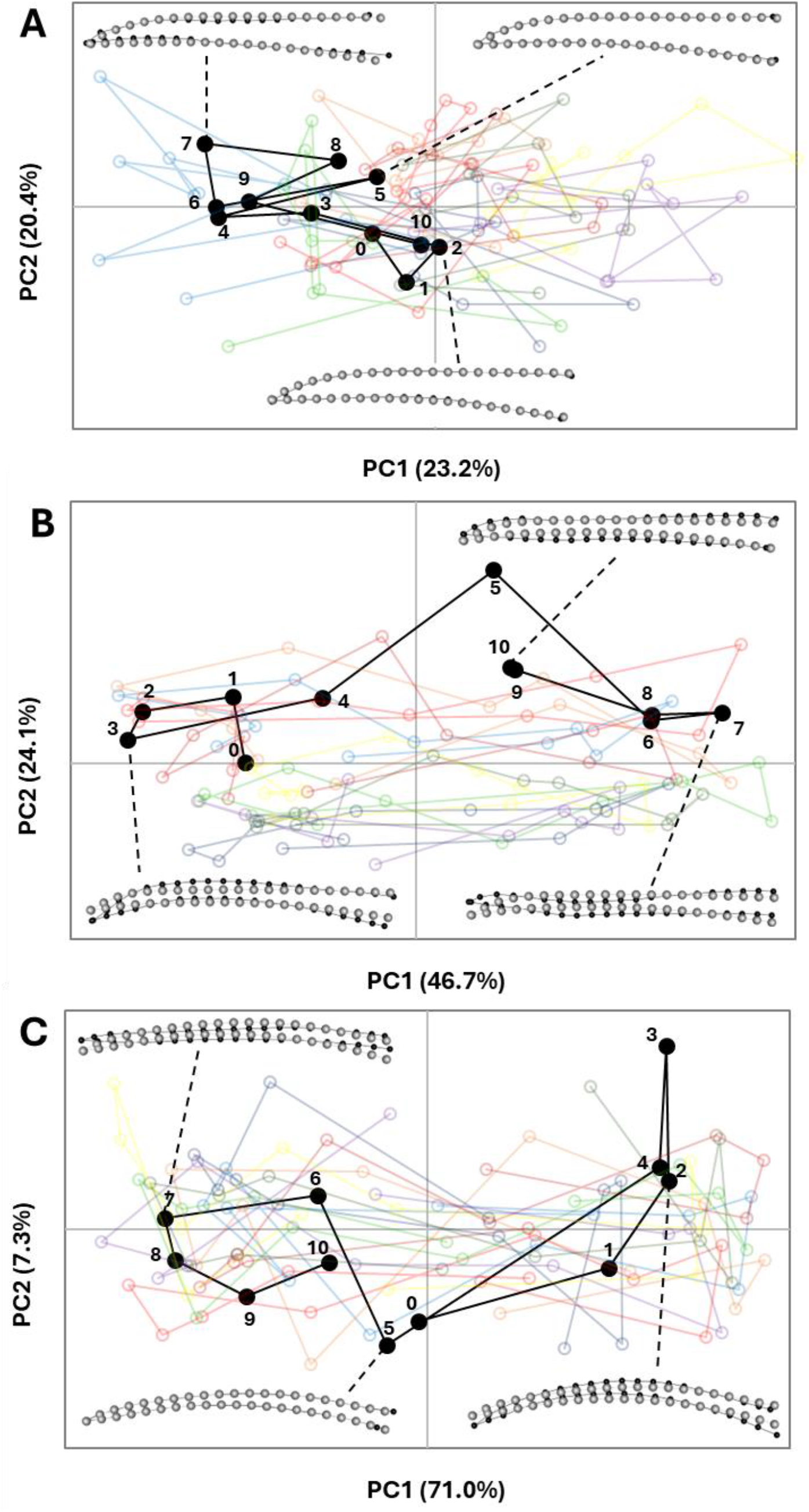
Shape changes in PC1-PC2 morphospace observed in the culmen of species with more aberrant overall bill shapes for A) *Florisuga mellivora*, B) *Myrmia micrura*, and C) *Phaethornis longirostris*, during lick cycles. In each graph, “Lick01” (black filled symbols, numbered sequentially) is shown for reference; the semi-transparent open symbols and traces in other colours are those of the other nine lick cycles (Lick02-10). The insets show representative shape changes (black dots and lines; relative the grey dots and lines of the consensus configuration, magnified by 7X) for selected frames during Lick 01 for each species.

### 3.2. Flexural rigidity and rhynchokinetic bill movements

Dorsoventral flexural rigidity patterns through the laterally positioned maxillary processes of the premaxillae often differed from patterns observed through the dorsally positioned frontal processes of the premaxillae. We identified three regions with distinct flexural rigidity profiles (figure S11): region 1, which is near the bill base and includes the nasal cavity; region 2, which constitutes most of the bill length; and region 3, where the lateral (ventral sensu Zusi [17]) and dorsal bars fuse; these regions roughly correspond to the nasal, intermediate, and symphysial parts of the prepalatal upper jaw (see Fig. 5 in [17]). Within region 2 (the “bending zone” identified from high-speed videography), both the lateral and dorsal bars of the upper bill exhibit substantially lower flexural rigidity values than they generally do in the proximal and distal rigid zones (regions 1 and 3). When the flexural trends through the upper jaw are viewed holistically, the bending zone exhibits substantially lower average dorsoventral flexural rigidity values than either the proximal or distal rigid zones, and this is consistent among species (figures S11, S12). These patterns of deformation and landmark displacement during the licking cycle seem to reflect two different forms of motion. On the one hand, the high GPA total variances reflect greater ranges (and speeds) of relative displacements of end members along the vertical (Y) axis (orthogonal to the length of the bill), as observed at the distal bill tip. On the other hand, the pattern of variance at the base, and to some extent the mid-region of the bill, are more likely a consequence of out-of-plane, lateral expansion and contraction of the bill, drawing the culmen and tomium dorsoventrally in those regions. Thus, in these regions, the observed variance and deformations are indicative of a dorsoventral bending zone (figure 5), likely facilitated by flattening of the cross-sectional arch of the culmen, which would otherwise resist dorsoventral bending. In most taxa, there is a clear peak in dorsoventral flexural rigidity, on either side of which much of the upper bill movement (i.e., “bending”) appears to be concentrated (figure 5). Naturally this varies among taxa; for example, in *Florisuga mellivora* (figure 5d), the upper bill movement is diffuse and does not distinctly correspond to the point of minimum dorsoventral flexural rigidity. In other taxa, such as *Amazilis amazilia* (figure 5a), *Calypte anna* (figure 5b) and *Phaethornis longirostris* (figure 5e), there appears to be an additional, more cranially positioned local minimum in dorsoventral flexural rigidity, resulting in an additional bending zone corresponding to high degrees of middle and cranial upper bill motion.

**Figure 5.**
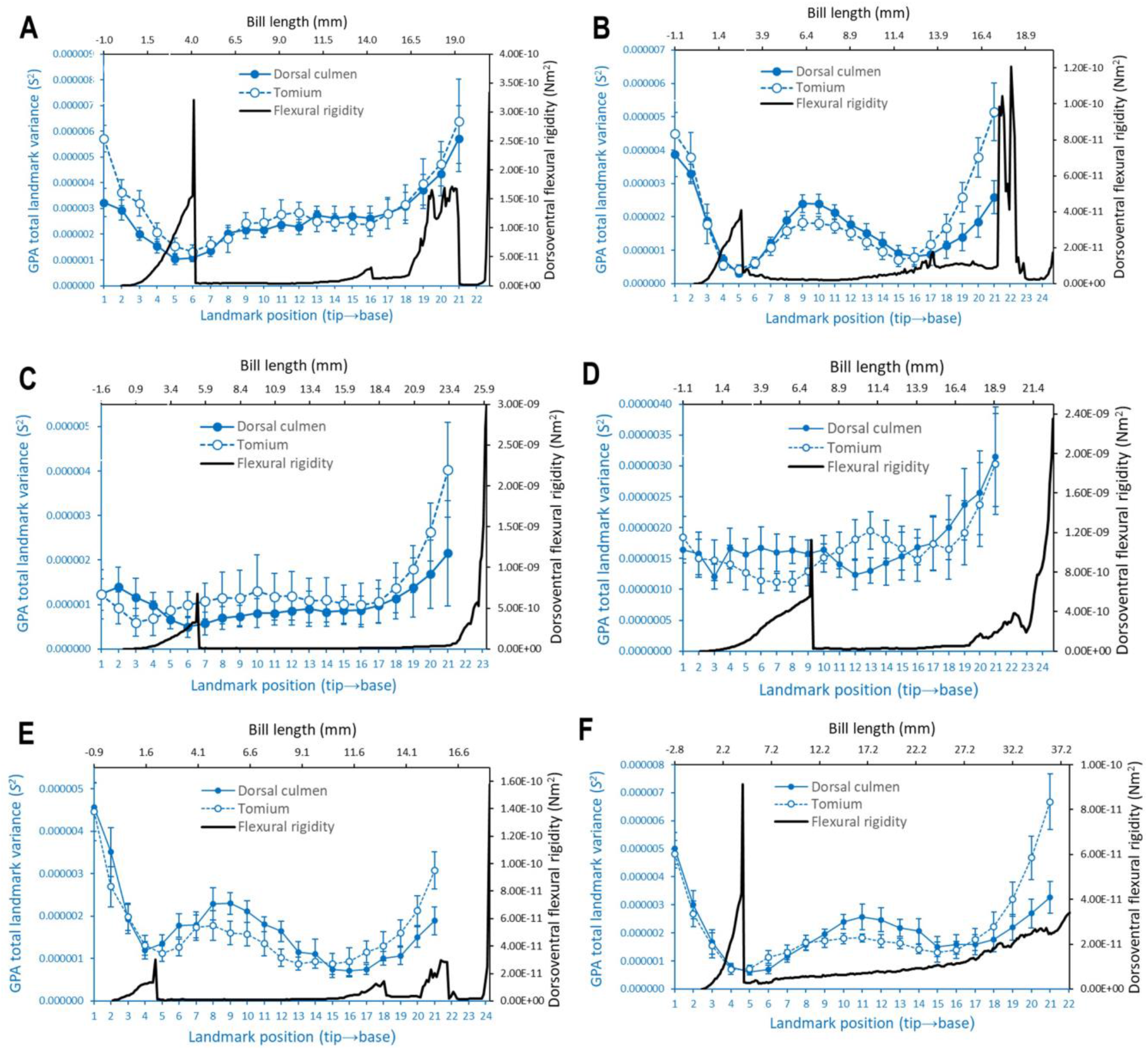
Mean ± 95% CI GPA total landmark variances (S^2^) computed over 11 sequential frames, across *n* = 10 lick cycles for each subject (blue symbols and lines along corresponding x [bottom] and y [left] axes): A) *Amazilis amazilia*, B) *Calypte anna*, C) *Chalybura buffonii*, D) *Florisuga mellivora*, E) *Myrmia micrura*, and F) *Phaethornis longirostris*. The blue filled symbols and solid lines correspond to the dorsal culmen and the open symbols and dashed lines correspond to the tomium. The dorsoventral flexural rigidity traces (black, along corresponding x [top] and y [right] axes) of representative specimens of each species from BendCT are superimposed to compare zones of relative structural rigidity to zones of relative kinesis along the lengths of the bills, from rostral tip to cranial base (peaks in flexural rigidity correspond to the point of fusion between the premaxillae rostrally and the craniofacial hinge [cranially]). The top horizontal (flexural rigidity bill length) axis starts at negative values accounting for the keratinous rostral tip of the culmen (from the CT scans, see Methods); similarly, the bottom horizontal (bill landmark) axis extends beyond 21 as it corresponds to the bony nasofrontal hinge.

**Figure 6.**
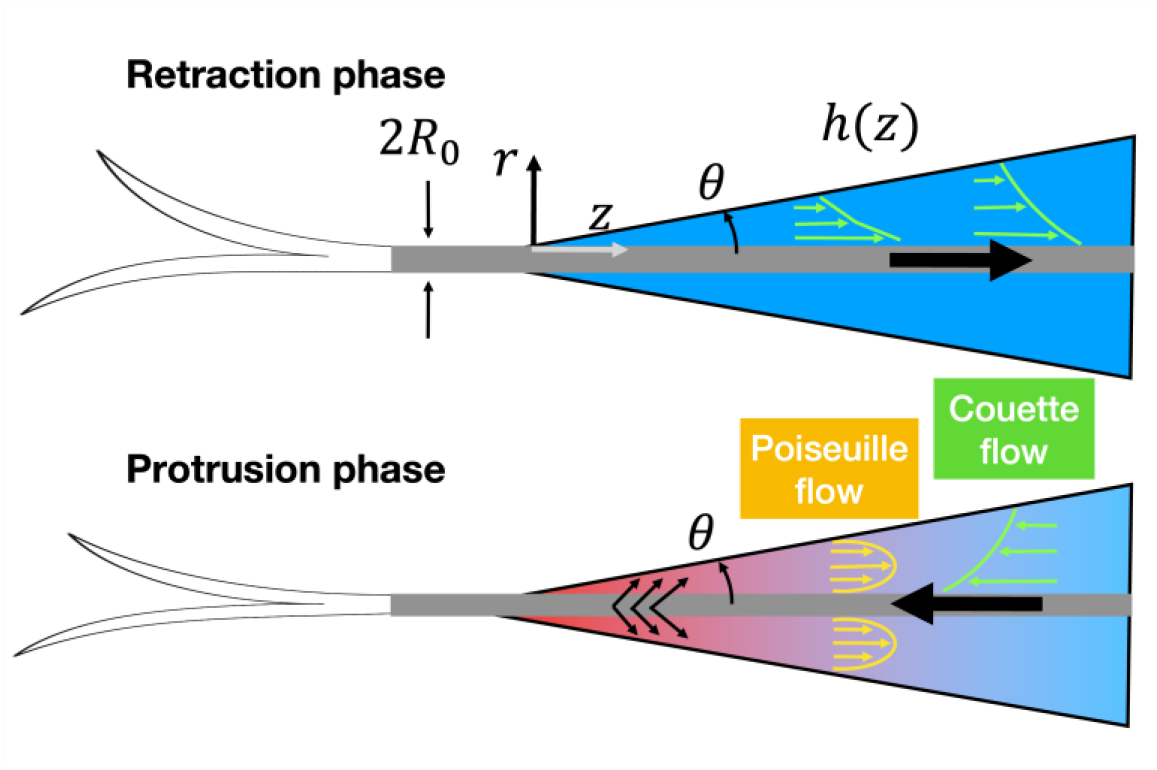
Two schematics showing fluid flows within the bill. The top panel illustrates the retraction phase, where the tongue is pulled back into the bill, creating a pure shear-driven fluid motion known as Couette flow. The bottom panel illustrates the protrusion phase, while the bill wrings the tongue, nectar is squeezed into the oral cavity near the tip while the tongue moves outward. The compression at the bill tip generates a high-pressure zone (indicated in red) which diminishes progressively along the tongue’s axis, transitioning to a low-pressure area (shown in blue). This axial pressure gradient gives rise to a pressure-driven flow (Poiseuille flow) that operates concurrently with the shear-driven Couette flow. The combined fluid mechanics during both phases facilitate the efficient transport of nectar.

### 3.3. Fluid analysis

There are two distinct motions involved in hummingbird drinking: angle change of the bill walls in relation to the tongue, and reciprocating tongue motion. First, change in the angle of the bill will affect the cross-sectional area of the oral cavity. When separation between the bill tips is large and separation at the bill base becomes small, the overall bill angle becomes smaller during the protrusion phase. Inversely, bill angle increases during the retraction phase. Such an angle change could modify the pressure gradient along the long axis of the bill. When the angle becomes smaller, the pressure on the bill tip will gradually increase, thereby pushing fluid towards the bill base. Thus, angle change between the bill walls and the tongue can contribute to the amount and duration of drinking.

Second, repeated movement of the tongue in and out of the bill can generate a shear flow within the bill, to transport nectar along the bill’s long axis. However, the cyclic motion of the tongue by itself would not create a unidirectional flow. To break the symmetry, the bill motion should be coordinated. The squeezing of the tongue by the bill tips wrings the nectar out of the tongue grooves. This motion also results in increasing the bill angle as well as inducing a nectar source inside the oral cavity.

To describe fluid motion inside the bill, let *v*_*z*_(*r, z*) denote the axial velocity of the fluid within the bill at a radial position *r* and along the axial position *z*. The velocity profile inside the bill can be described by

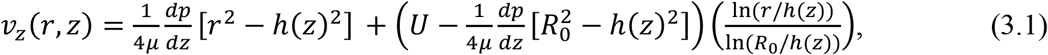

Where *μ* is the dynamic viscosity of the fluid, *dp/dz* is the pressure gradient along the axial direction, *h*(*z*) is the radius of the fluid column at position *z, R*_0_ is the radius of the tongue, and *U* is the velocity of the tongue’s linear motion.

The terms with *dp/dz* represents the flow contribution from the axial pressure gradient and the term with *U* represents the shear flow due to the tongue’s linear motion. The volumetric flow rate *Q*_0_ can then be obtained by integrating the axial velocity over the radial cross-section, from the tongue *r* = *R*_0_ to the boundary *r* = *h*(*z*), giving

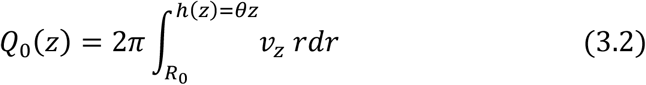

This integral will provide the total flux for a given axial position z, assuming axial symmetry of the flow and constant *dp/dz*.

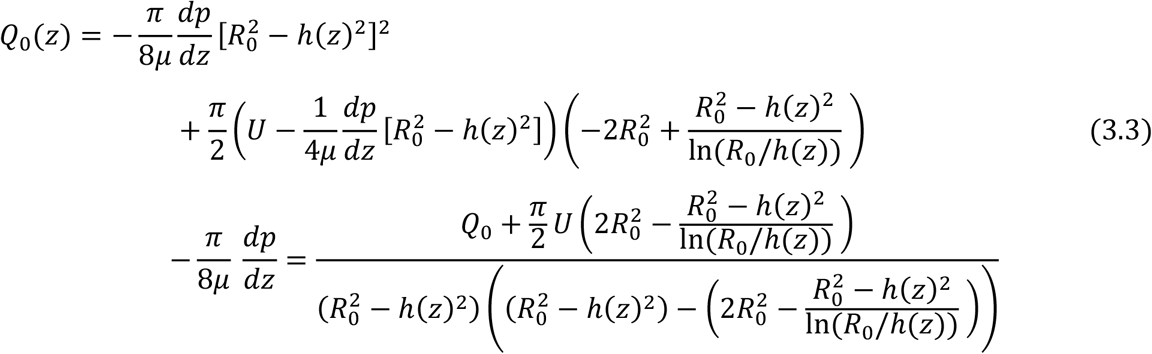

The volumetric flow rate *Q* within the bill must remain invariant with respect to the axial position *z*, satisfying the conservation of mass in the system. Therefore, regardless of the position along the axis of the bill, we assert that *Q* = *Q*_0_, where *Q*_0_ is the volumetric flow rate at a reference cross-section.

To estimate the energy cost associated with lapping, we evaluate the wall shear stress on the tongue. This viscous-resisting force is computed by the following relationship

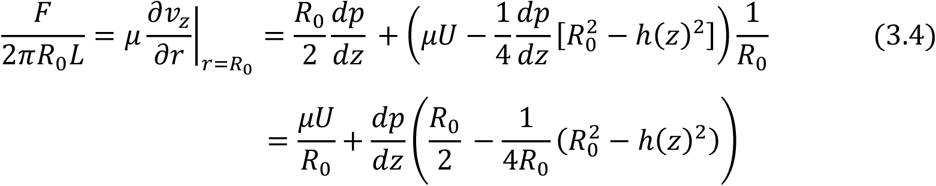

Where τ is the wall shear stress and *L* is the tongue length inside the bill.

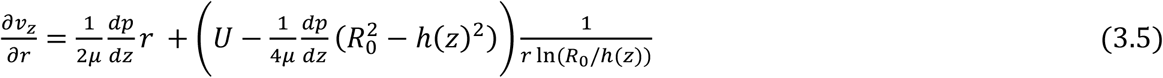

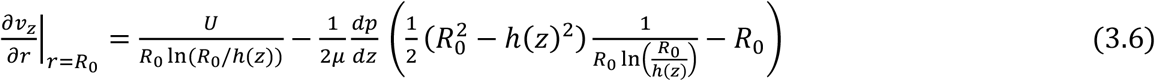

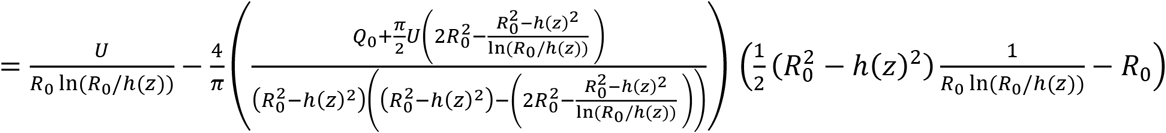

In the asymptotic limit *R*_*o*_ ≪ *h*(*z*), the shear stress on the tongue can be approximated. The force per unit length due to shear stress simplifies to

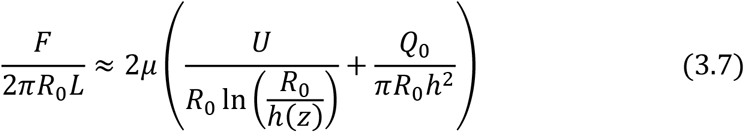

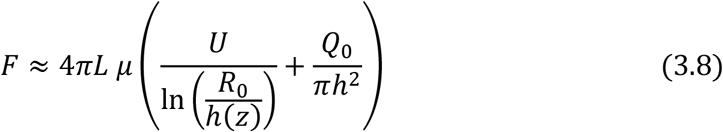

Here, the flow rate *Q*_0_ can be approximated as 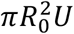, which is based on the assumption that hummingbirds efficiently scrape out all nectar from tongue grooves during extraction. Hence, the second term depends on 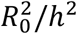, which is slowly varying compared to the logarithmic first term ln(*R*_0_/*h*). This implies that the viscous resistance faced by the tongue is significantly affected by the first shear-flow term, particularly by the velocity *U* and bill geometry.

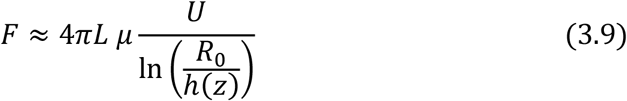

This indicates that increasing the gap between the tongue and the beak, effectively increasing *h*(*z*) relative to *R*_0_, reduces the resisting force due to shear stress. If the bill has a blunt-tip configuration, characterised by a rapid expansion in the height *h* of the fluid column, the frictional force experienced by the tongue can be substantially reduced. Such a reduction in resisting force is energetically favourable as it potentially lowers the energy required for nectar feeding.

## 4. Discussion

### 4.1. Rhynchokinesis in hummingbirds

Hummingbird maxillae are inherently flexible because both the dorsally positioned bar (bone along the top of the bill, corresponding to the fused frontal processes of the premaxillae) and ventro-laterally positioned bars (bones on the sides of the upper bill, corresponding to the maxillary processes of the premaxillae) are thin, elongated, and flattened [17,45]. Nitzsch [50] proposed a form of rhynchokinesis in hummingbirds as an adaptation for nectar feeding, consisting of extensive flexion zones along the dorsal and ventral aspects of the upper jaw (see [14]). However, Zusi [16] found no osteological evidence to support any distal rhynchokinetic capacity in hummingbirds, leaving the prospect of rhynchokinesis in hummingbirds questionable. However, videos later presented by Rico-Guevara and Rubega [48] led Zusi [17] to conclude later that distal rhynchokinesis could occur in hummingbirds, possibly through sliding of the ventrolateral bone bars relative to the dorsal bone bar, and intrinsic bending of the ventrolateral bars.

Although the question of hummingbird maxillary bending has not been directly broached in the years since, Rico-Guevara et al. [49] showed phase-shifted bill opening where the jaws near the bill tip and near the bill base can be separated at different points throughout the licking cycle, while maintaining the middle portion of the bill mostly shut during the licking cycle. Such a mechanism could work via bending or rotation of the mandible, ultimately lowering the tip of the lower jaw, or by bending of the upper jaw to deflect the tip upwards, or by combinations of those mechanisms. Given their extensive mandibular flexibility [15,27,28], we expected to find support for mandibular bending to some extent. Instead, our results illustrate clear evidence of upper bill bending in hummingbirds—the first time that maxillary bending near the bill tip has been quantitatively described in this group. Following Zusi’s [16] classification of bill bending, hummingbirds would perform either amphikinesis or distal to extensive rhynchokinesis (for those species previously reported to be prokinetic [17]), or double rhynchokinesis (for those species previously reported to be proximally rhynchokinetic [17]). However, the exact functional categories to which particular hummingbird taxa are best assigned may vary, and clarification must await further anatomical and morphofunctional studies of a wider range of hummingbird species.

By combining high-speed video of bill bending, coupled with a microstructural analysis of bone rigidity, we found that while the mandible (lower jaw) remains relatively in place throughout the licking cycle during nectar feeding, the distal portion—and, to some extent, the middle portion— of the maxilla (upper jaw) bends upwards. With the possible exception of *F. mellivora*, all taxa studied showed discernible rostral, and in some cases middle and cranial, bending zones (where most of the upper bill movement was concentrated) that are punctuated by points of substantially greater dorsoventral flexural rigidity. We loosely interpret these points as “hinges,” beyond which areas of much lower dorsoventral flexural rigidity enable the culmen to bend. The data presented here verify the presence of a region of relatively low dorsoventral flexural rigidity in the upper jaws of several hummingbird taxa. As predicted by high-speed videography, these regions of low flexural rigidity roughly correspond in their position to the region of dorsally directed upper jaw bending and affirm the existence of proximal and distal rigid zones. The dorsal bending of the hummingbird upper jaw adds to the simultaneous ventral and lateral deflection of the lower jaws, measured in *Archilochus colubris* during insect hawking events [15], as another means by which hummingbirds achieve enhanced gape sizes. Future work will seek to characterise the mechanism by which this lateral and ventral mandibular deflection occurs; it seems possible that regions of decreased flexural rigidity underlie the ability of hummingbird lower jaws to deflect in this manner as well.

### 4.2. Implications of bill bending for intraoral fluid transport

To maximise nectar uptake efficiency, offloading as much nectar as possible from the tongue is paramount for maintaining the maximum loading capacity across consecutive licks [45]. After wringing the tongue to unload the nectar inside the bill, this volume of liquid must be transported towards the throat to be swallowed. In hummingbirds, such intraoral transport must be coordinated with tongue squeezing at the bill tip at rates exceeding several times per second. A bending point in the middle-to-distal region of the bill tip has the potential to coordinate these two processes, by allowing the tip to open and close (to provide nectar on- and off-loading) while keeping the middle-bill region tightly closed to provide a fairly constant internal diameter for the “tongue squeegee” to work properly, wherein the flaps at the base of the tongue can maintain close contact with the intraoral cavity walls to move the nectar proximally. Previously, Rico-Guevara et al. [49] described intraoral transport mechanisms during nectar drinking in hummingbirds (distal wringing, tongue raking, and basal expansion). Here, we present the functional basis of the maxillary flexibility that allows those mechanisms to occur and be synchronised at high rates. Hummingbirds drink nectar by reciprocating their tongues up to 17 times per second [72]. Bill and tongue movements must therefore be well coordinated at this fast pace, as the bill tips must close immediately after the tongue ends its retraction and before it starts its protraction to maximise the amount of nectar retained inside the bill (distal wringing sensu Rico-Guevara et al. [49]). Nearly closing the bill tips while extruding the tongue through them creates a diminished volume of the distal oral cavity (reduced cross-sectional area) and a sharp angle of the internal bill walls as they contact the tongue (Figure 1 in [45]). This operation is vital not only to move nectar from the tongue to the bill, but also to clean and compress the tongue in order to collect as much nectar as possible in the following lick [46]. Distal rhynchokinesis allows for the bill tips to be open during the tongue retraction phase and to be kept mostly closed during the extension phase while preventing jaw separation around the middle portion of the bill, which ultimately facilitates the tongue base raking the nectar [49]. A hummingbird’s tongue base is analogous to a stick squeegee used to clean paintball gun barrels, which consists of a rod with a rubber disc at one end that swivels in such a way that when inserted it does not displace the fluid, and when retracted forms a seal against the walls of the barrel pulling fluid in the same direction. The equivalent to the swivelling rubber disc component in hummingbirds is the tongue base with folding flaps (i.e., tongue wings).

Rhynchokinesis near the bill tip allows for close contact between the tongue base flaps and the walls of the intraoral cavity along the middle portion of the bill, while allowing the next aliquot of nectar to be drawn into the bill cavity by the loaded tongue through the open tips. Thus, distal rhynchokinesis enables expansion of the intraoral space near the bill base to occur in a phase-shifted manner, in such a way that, at the same time that nectar is released into the oral cavity by the nearly closed bill tips, the bill base is open, increasing intraoral capacity for nectar to flow into an enlarged space towards the throat. We propose that this type of rhynchokinesis near the bill tip improves intraoral nectar transport and therefore feeding efficiency. There is variation across species in the degree of bill bending (e.g., figure S13), which warrants further studies on the potential relationships with their ecology and other selective pressures on bill performance (discussed below). For instance, hummingbirds with short and straight, stout bills might not benefit as much from the advantages of the coordination of intraoral transport mechanisms described below, but on the other hand, might not need to, given the comparatively limited transport distances for nectar off-loading and transport to the throat.

### 4.3. Potential trade-offs and constraints on the evolution of bill bending in hummingbirds

The hummingbird bill can work as a tight unit with clockwork precision coordinating with rapid tongue movements many times a second [31,49]. However, it can also be decoupled as when the gape is expanded by mandibular bowing during displays or arthropod hunting [15,27,28]. Although hummingbirds exhibit the capacity to extensively bend their lower jaws during aerial insectivory [15,27], while drinking nectar hummingbirds hold their mandibles relatively still (e.g., Movie S1), and appear to use them as a base of support upon which to bend their maxillae while drinking nectar. Since flycatching bill manoeuvrers rely on rapid closure by elastic instability of the mandibular bending [27], it is possible that maxillary bending and passive rapid unbending could also aid during arthropod foraging.

Hummingbird bills are also a key structure to access nectar resources beyond their interaction with flowers. Since a given flower could be depleted by many individuals and often from several species competing in the same area, hummingbirds face interference competition, using their bills as weapons [31,73]. Hence, a bill adaptation expected to enhance fighting proficiency is increased structural soundness to withstand impacts. This would select for overall bill robustness, as more solidly built structures better withstand potentially damaging bending forces and would also transfer larger stabbing and biting forces to the bill tip. Therefore, the maxillary bending and structural flexibility described here could face a tradeoff between improving nectar drinking by allowing for opening of the bill tips while the middle portion of the beak remains tightly closed (enhancing intraoral transport) and improving fighting performance when the bill is used to fence off rivals during territorial quarrels for floral resources. Under the axial load likely to be exerted when the bill is employed as a stabbing weapon, such structural flexibility is likely to cause buckling and even failure. Thus, increased flexural rigidity will improve stabbing performance by diminishing bill bending, but at the same time it will negatively affect nectar drinking efficiency. We expect that the patterns of flexural rigidity (and therefore bending) of hummingbird bills vary intra- and inter-specifically. Adult males are more frequently reported defending floral patches than females and juveniles [74–77], consequently, we expect the bills of males (or individuals that engage in fights more often) to present higher dorsoventral flexural rigidity when compared to females and juveniles. As shown here, there is great variation in bill length (table S10) and size-controlled shape (figure S10) across hummingbird species (with even more variation across the family, e.g., [78]). Flexural rigidity will be affected by such variation in bill architecture, in addition to potential changes in material properties occurring in response to particular selective pressures such as the use of bills as intrasexually selected weapons [79], and as tools for visual displays (e.g., differences in melanisation across the bill length, Fig. 3 in [49]). For example, a bill trait that greatly affects structural properties is curvature because, when loaded axially, elongated structures are mechanically more resistant to buckling if they are straight [80,81]. As such, bending is disadvantageous for a stabbing weapon, as less force is applied at the tip and subsequently less damage can be done to an opponent. In hummingbirds, straighter bills transmit more force without bending, and pointier bills transform that force into perforation capacity [73]. Therefore, we expect differences arising from pressures of their bills used as weapons, as well as intrinsic differences in bending given their bill architecture (e.g., maxillary thickness and curvature, figure S10).

The extent to which hummingbirds can perform rhynchokinesis near the bill tip depends on the overall shape and size of the bill. We found that the degree of bill bending is greater in bills with relatively more curved distal, and narrower proximal, shapes for their size, and lower in bills characterised by relatively straighter distal, and deeper proximal, shapes for their size (electronic supplementary materials, S1.2; figure S13). However, a more comprehensive assessment, including a greater diversity of species in a phylogenetic comparative context (and taking advantage of recent 3D characterization tools, e.g., [78]), is certainly warranted to substantiate this trend.

### 4.4. Conclusions

We argue that maxillary bending in hummingbirds, arguably the most specialised avian nectarivores, is a functional innovation that improves feeding efficiency and may constitute an important aspect of morphological diversification within plant-hummingbird coevolutionary systems. Hummingbirds can open their bill tips and separate the base of the jaws while maintaining a tightly closed middle section of the bill, which enhances the efficiency of nectar extraction because it allows nectar to be taken up at the bill tips (when offloaded from the tongue) and intraorally transported to the throat (through a tight bill tube) simultaneously. This enables hummingbirds to exploit longer and narrower flowers than they would be able to if their bills were inflexible and if they had to open the entire bill to separate their bill tips. The particularly elongate and narrow corollas that can evolve under this scenario, wherein bill and flower shape match, would benefit both birds and plants [82], perhaps evolutionarily filtering less efficient pollinators.

## Acknowledgements

Dedicated to the memory of functional morphologist and hummingbird anatomy authority, Richard L. Zusi. Thanks to Doug Altshuler and Chris Clark for the use of their footage of captive hummingbird feeding for preliminary analyses (which were not included in this published version) and for providing specimens for CT scanning. Edward Hurme and Jesse Joy assisted in the field and with tracking image sequences, Yiran (Melissa) Liu and Jason Fan participated in the tracking error analyses. Special thanks to Margaret Rubega for her encouragement and support of this work, and to Kurt Schwenk for helpful discussions along the way. Thanks to AnaMeli Fernandes for manuscript and figure formatting, as well as Alyssa Sargent, Felipe Garzón, and Sam Case for general revision. We thank Rick Prum and Kristof Zyskowski for access to specimens, and Jessie Maisano and Joshua van Houten for CT scanning assistance. For the purpose of open access, the authors have applied a Creative Commons Attribution (CC BY) licence to any Author Accepted Manuscript version arising.

## Funding

A.R.-G. is supported by the Walt Halperin Endowed Professorship and the Washington Research Foundation as Distinguished Investigator. D.J.F. is supported by UKRI grant MR/X015130/1.

## Author contributions

ARG, DJF, and DS conceived the study; ARG and KJH obtained video footage in the field; ARG, KJH, and JEH tracked images; ARG and KJH performed the baseline kinematic analyses; DS performed the geometric morphometric analyses; DJF and JEH performed the BendCT flexural rigidity analyses. SJ performed the fluid analysis. All authors contributed to the writing and editing of the manuscript.

## Notes

### Competing Interest Statement

The authors have declared no competing interest.

