## Supplemental material for "Upper bill bending as an adaptation for nectar feeding in hummingbirds"

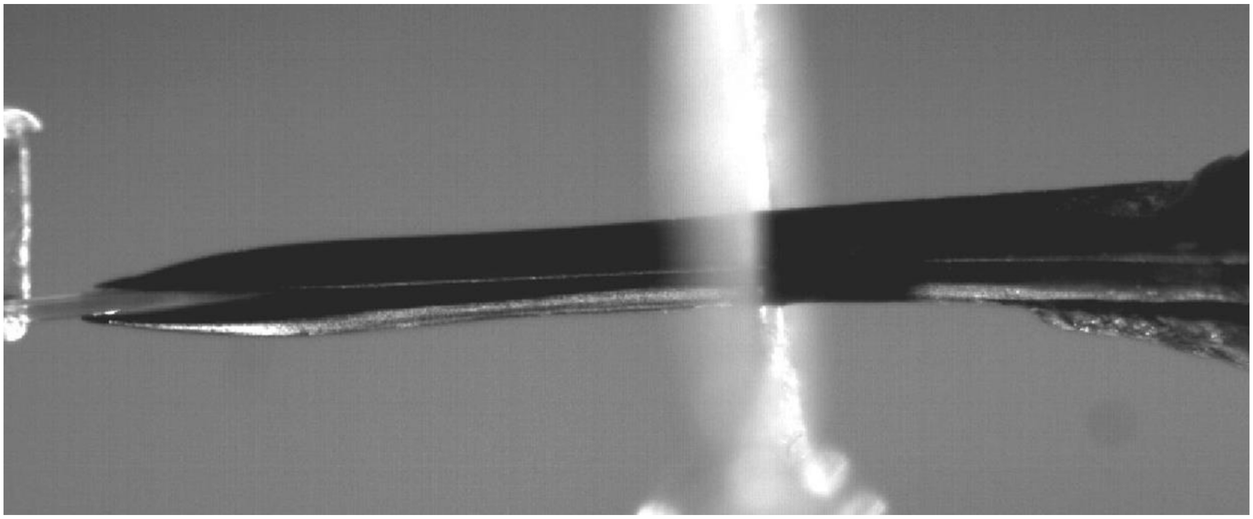

**Movie S1. Lateral view of a hummingbird drinking nectar.** High-speed video (1000 fps) of an Anna's Hummingbird (*Calypte anna*), the base of the bill is on the right and the nectar reservoir on the left. The diffuse shadow in the middle is the flat, flower-shaped guide that served to control the hummingbird positioning.

### Supplementary Material S1

#### S1.1. Analysis of Tracking Error and Repeatability

Because three different researchers ultimately participated in tracking the final dataset of 660 images, we investigated the potential effects of tracking error in two ways. For this analysis we used one randomly-selected lick cycle ("Lick02") of the *Calypte anna* subject, because it possesses what we feel is the most generalized bill shape among the species we studied. Five different participants tracked each frame three times, in random order to eliminate any sequence bias ($n$ = 5 participants × 3 repetitions × 11 sequential frames = 165). Therefore, these datasets contain both within- and among-participant tracking error, which we did not attempt to tease apart. We conducted a geometric morphometric analysis of culmen shape changes during the lick cycle (as described above), followed by a Procrustes ANOVA to test the significance of any differences among sequential frames, despite intra- and inter-participant sources of variance (adapted from approaches by [1,2]). For this analysis, "participant" was included as a factor, with "repetition" nested within participant. Secondly, we performed an analysis of landmark precision (adapted from [3,4]) to ensure that we could distinguish upper bill deformation signal from noise (e.g., within- and among-participant tracking error) during a lick cycle. We performed GPA (as described above) and generated total landmark variances among the 15 repetitions of each frame image separately, resulting in a GPA total landmark error variance value ($S^2$) for each landmark, for each frame. We then computed the mean ± 95% CI total landmark variances across all ($n$ = 11) frame images of the sequence, collectively. Our assumption was that variation within repeated frame images should be less than that among sequential frame images, but given the added among-participant variance, this approach provides a very conservative assessment of the signal: noise ratio.

The degree of shape change in the culmen among frames during a licking cycle followed a specific trajectory through shape morphospace (mostly along PC1, which accounted for most of the variance), that was repeatable despite considerable tracking error (mostly along PC2) within and among participants (figure S1). Despite the significant differences among participants (Procrustes [nested] ANOVA; $F_{10,10}$ = 9.01, $P$ = 0.0129; table S1), repetitions within participants did not vary significantly ($F_{10,140}$ = 0.366, $P$ = 0.544), and bill shape changed significantly across sequential frames ($F_{10,140}$ = 31.88, $P$ = 0.0001; figure S2), with variation among frames accounting for most of the variance in the model ($R^2$ = 0.63). The predominant axis of shape variation was characterized by a relative dorsoventral narrowing of the middle of the bill, combined with a relatively upturned rostral third of the bill along the negative end of PC1, compared to a relative dorsoventral widening of the middle of the bill, combined with a relatively decurved rostral third of the bill along the positive end of PC1. The shape variation observed along PC2 seemed to emanate mostly from intra- and inter-participant error (figure S2), characterized by a relative dilation (negative PC2) and relative compression (positive PC2) of the landmark configurations (figure S1).

#### S1.2. Effect of overall bill size and shape on range of bill bending

We performed a standard, 'static' geometric morphometric analysis of overall bill shape differences among species prior to the onset of any upper bill deformation. For this analysis we PCA of bill shape variation among species, using only the initial frames ("frame0") of each lick cycle for each species, to capture the bill in a relatively neutral position prior to the onset of the licking cycle. We followed this analysis with a Procrustes ANOVA, as described above, to examine the effects of species, centroid size, and their interaction on shape (using Type III Sums of Squares and Cross-products and set to 9,999 permutations of null model residuals). We then tested whether bill shape and size affect the range of bill bending observed for each species. We used, measured here as the GPA variance of the rostral-most dorsal culmen landmark ('lm 1') for each lick cycle as a measure of the range of deflection of the culmen tip, for which we used the GPA variance of the rostral-most dorsal culmen landmark ('lm 1') for each lick cycle. The GPA landmark variance was computed from 11 sequential frames of a lick cycle, then relativized to the minimum GPA variance observed among landmarks throughout the lick cycle. We used the lm function of base R [5] to run a general linear model testing the effects of static bill shape and log centroid size (independent variables) on log landmark 1 GPA variance (dependent variable), while accounting for variation among species (fixed factor). We used a univariate composite variable that encapsulated both bill size and shape in the form of shape "regression scores", obtained from the plotAllometry function of R package "geomorph" [6]. These regression scores are derived from the covariation between shape and the regression coefficients for log centroid size, and thereby reflect the shape changes predicted by the regression model, including residual variation in that direction of the regression vector for centroid size [6,7]. Deformation grids showing shapes associated with these regression scores were generated using the picknplot.shape function of R package geomorph [6].

The overall shape of the culmen (based on frames captured prior to initiating lick cycles) differed significantly among species (Procrustes ANOVA; $F_{5,48} = 424.0$, $P = 0.0001$; table S8; figure S10). There was also significant allometry in culmen shape variation (with respect to centroid size), such that larger bills were associated with a greater degree of overall curvature ($F_{1,48} = 7.93$, $P = 0.0001$; table S8; figure S10). These results are quantitatively similar even after removal of the large, and morphologically extreme, *Phaethornis*. We then asked whether the degree of bill bending depends on the underlying bill shape and size variation observed among species. We found that the range of rostral bill tip deflection, based on the relative GPA variance of the most rostral dorsal culmen landmark ("LM 1"), was significantly linearly related to bill size and shape ($P < 0.0001$; table S9), after accounting for significant variation among species ($P < 0.0001$; table S9; figure S13).

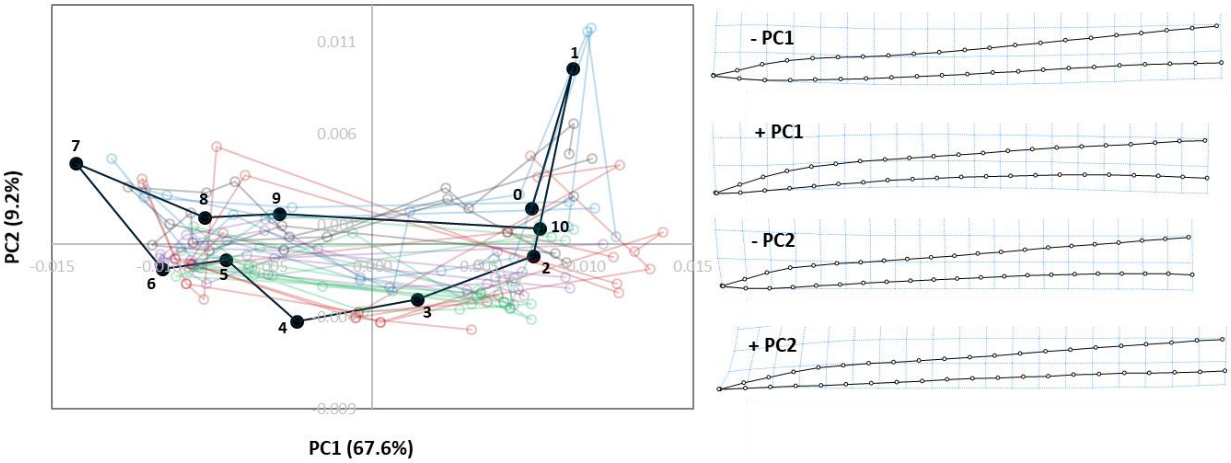


**Figure S1**. Shape changes observed in the culmen of *Calypte anna* in PC1-PC2 morphospace during a single lick cycle ("Lick02") tracked by five different participants, three times each (*n* =15). One arbitrarily selected repetition of one participant (black filled symbols and traces) is numbered sequentially for reference, and each semi-transparent set of open symbols and traces are those of other participants (each participant in a different colour). The deformation grids to the right of the graph show the ends of each PC axis.

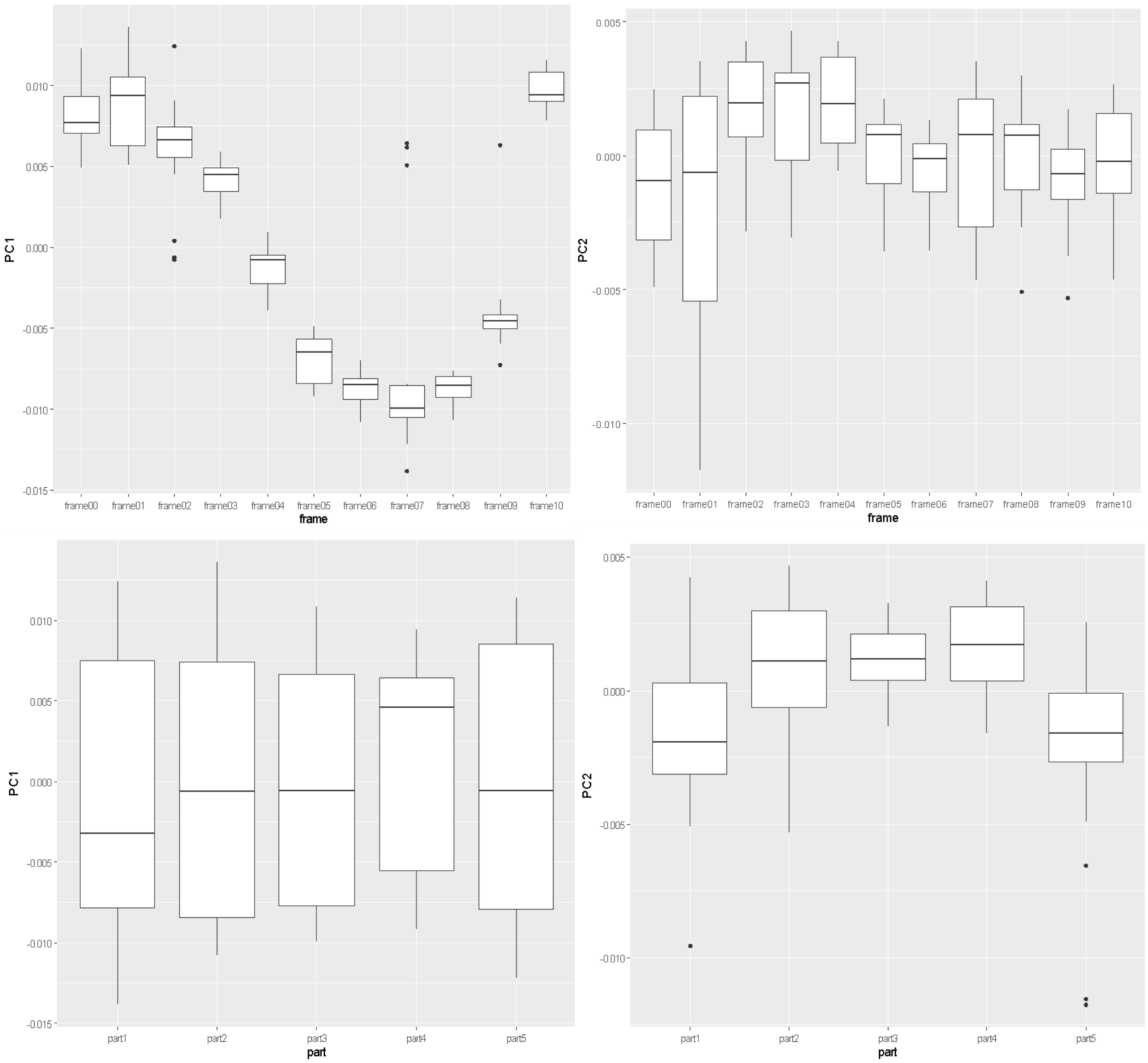


**Figure S2**. Series of boxplots showing variation among sequential bill landmark configurations, accounting for variation among repeated tracks of the same lick cycle (“Lick02”), along the PC axes (PC 1 left, PC 2 right) shown in figure S1 for *Calypte anna*, by frame (pooled over participants and repetitions; top row) and by participant (pooled over frames and repetitions, bottom row).

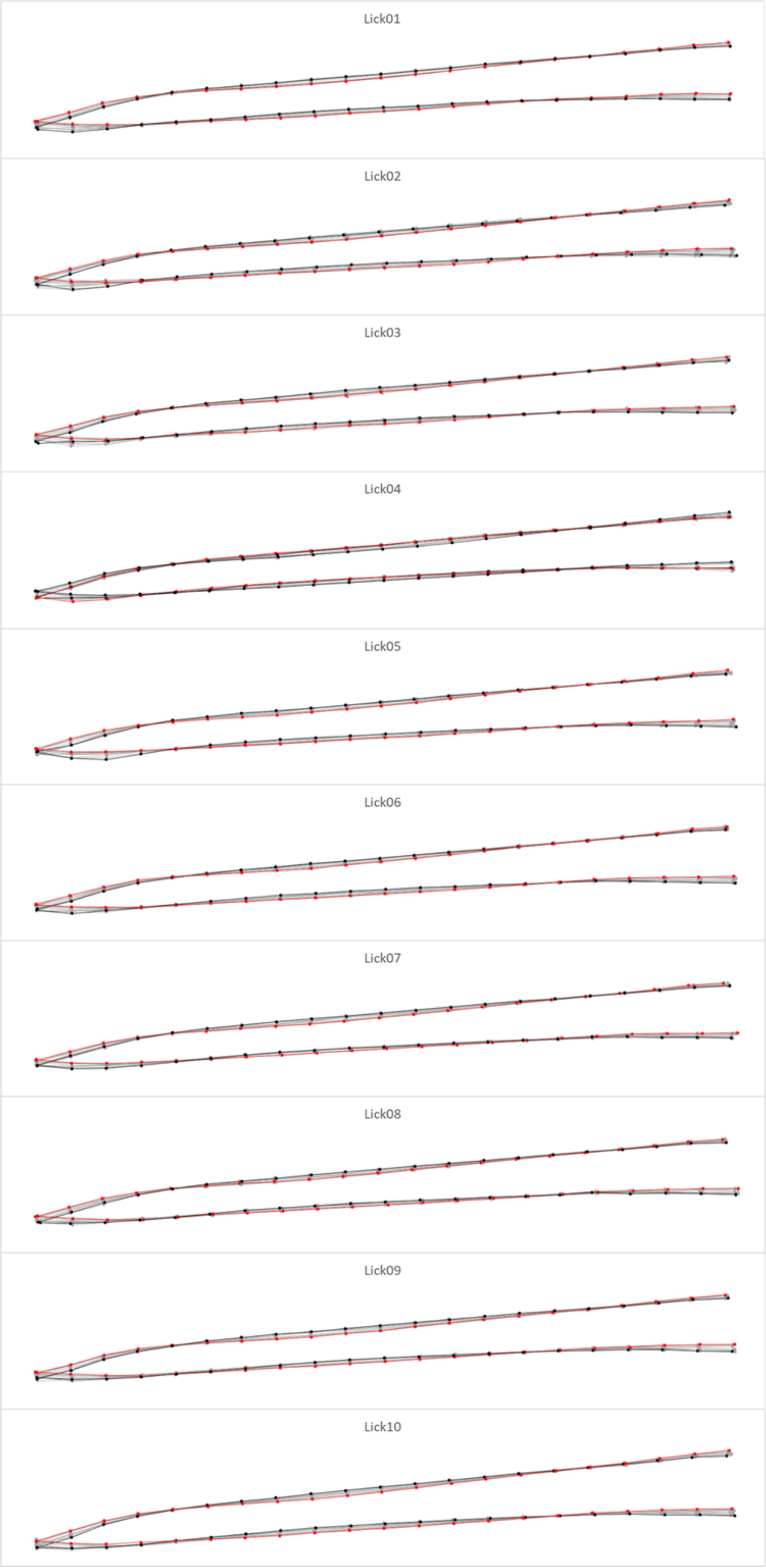


**Figure S3**. Alternative way of viewing shape changes during lick cycles. Aligned landmark configurations (by GPA; semilandmarks unslid) of 11 sequential frames for each of 10 *Calypte anna* lick cycles. Black and red outlines demarcate the two frames with the most dissimilar bill shapes during each lick cycle based on maximum pairwise Procrustes distances (all other frames are outlined in grey).

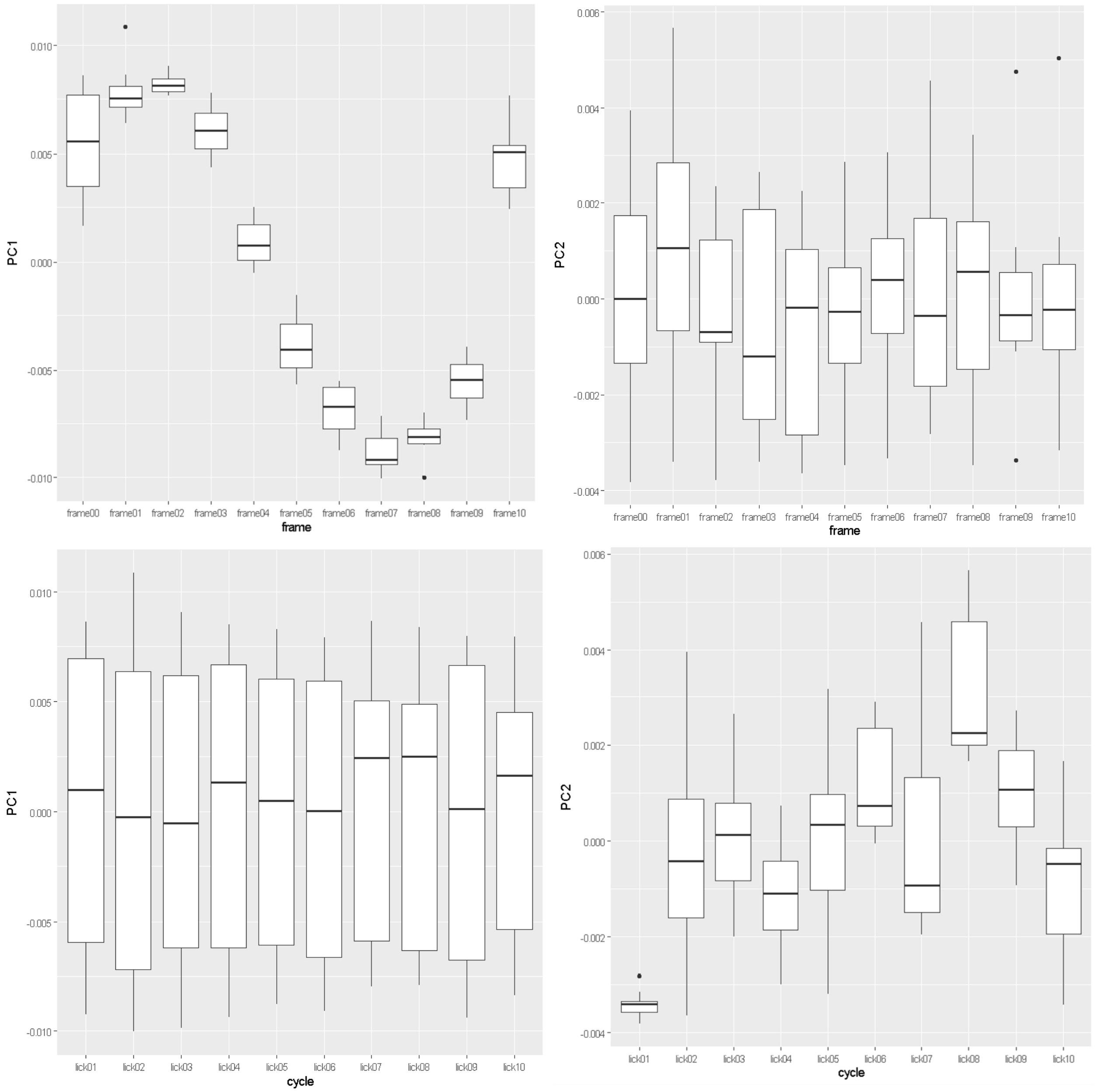


**Figure S4**. Primary sources of variation along the first two shape variables (PCs) that explain most of the variance in bill shape change for *Calypte anna*. Series of boxplots showing variation among sequential bill landmark configurations, accounting for variation among lick cycles, along the PC axes (PC 1 left, PC 2 right) shown in figure 3 for *Calypte anna*, by frame (pooled over lick cycles; top row) and by lick cycles (pooled over frames, bottom row).

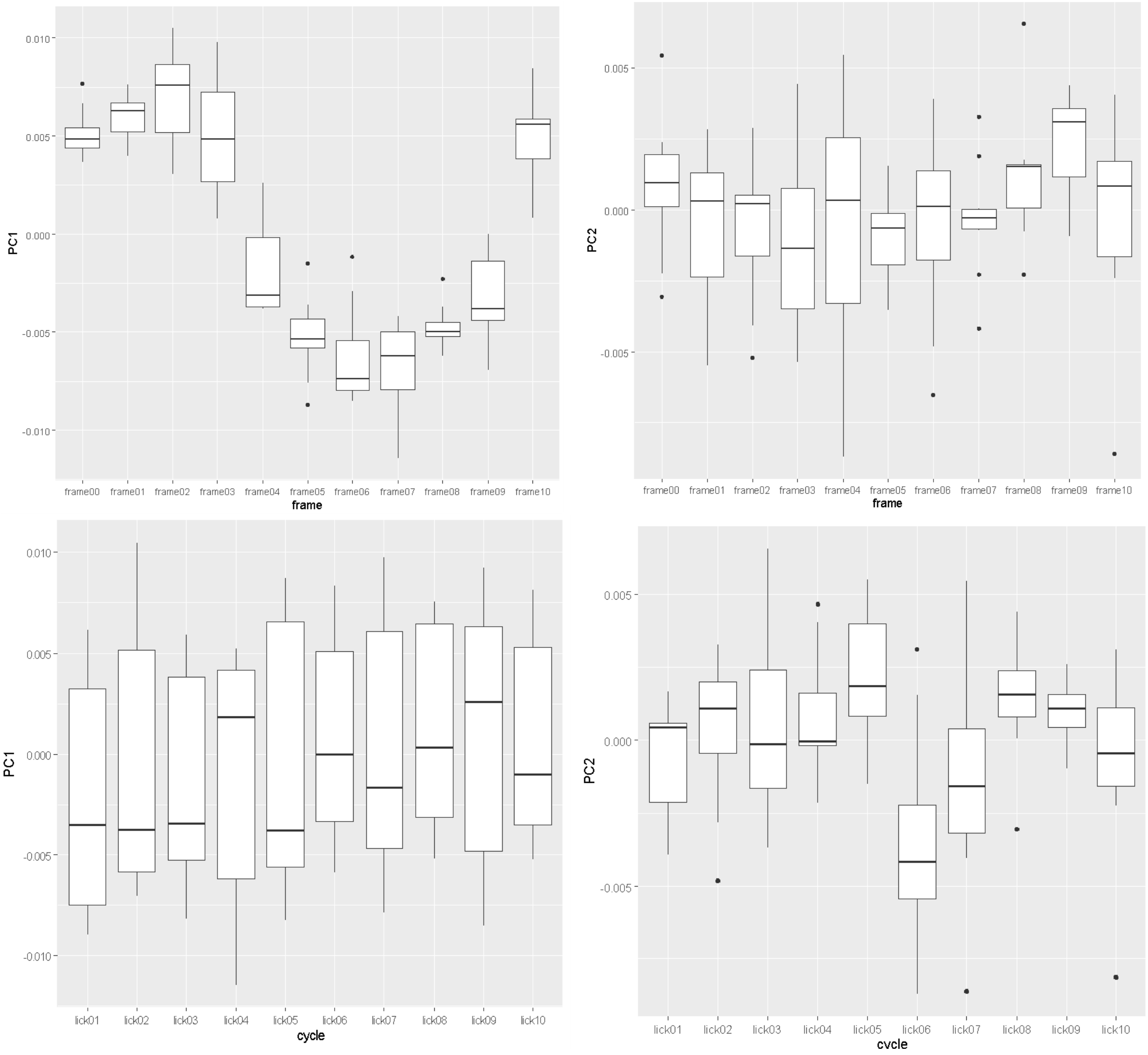


**Figure S5**. Primary sources of variation along the first two shape variables (PCs) that explain most of the variance in bill shape change for *Amazilis amazilia*. Series of boxplots showing variation among sequential bill landmark configurations, accounting for variation among lick cycles, along the PC axes (PC 1 left, PC 2 right) shown in figure 3 for *Amazilis amazilia*, by frame (pooled over lick cycles; top row) and by lick cycles (pooled over frames, bottom row).

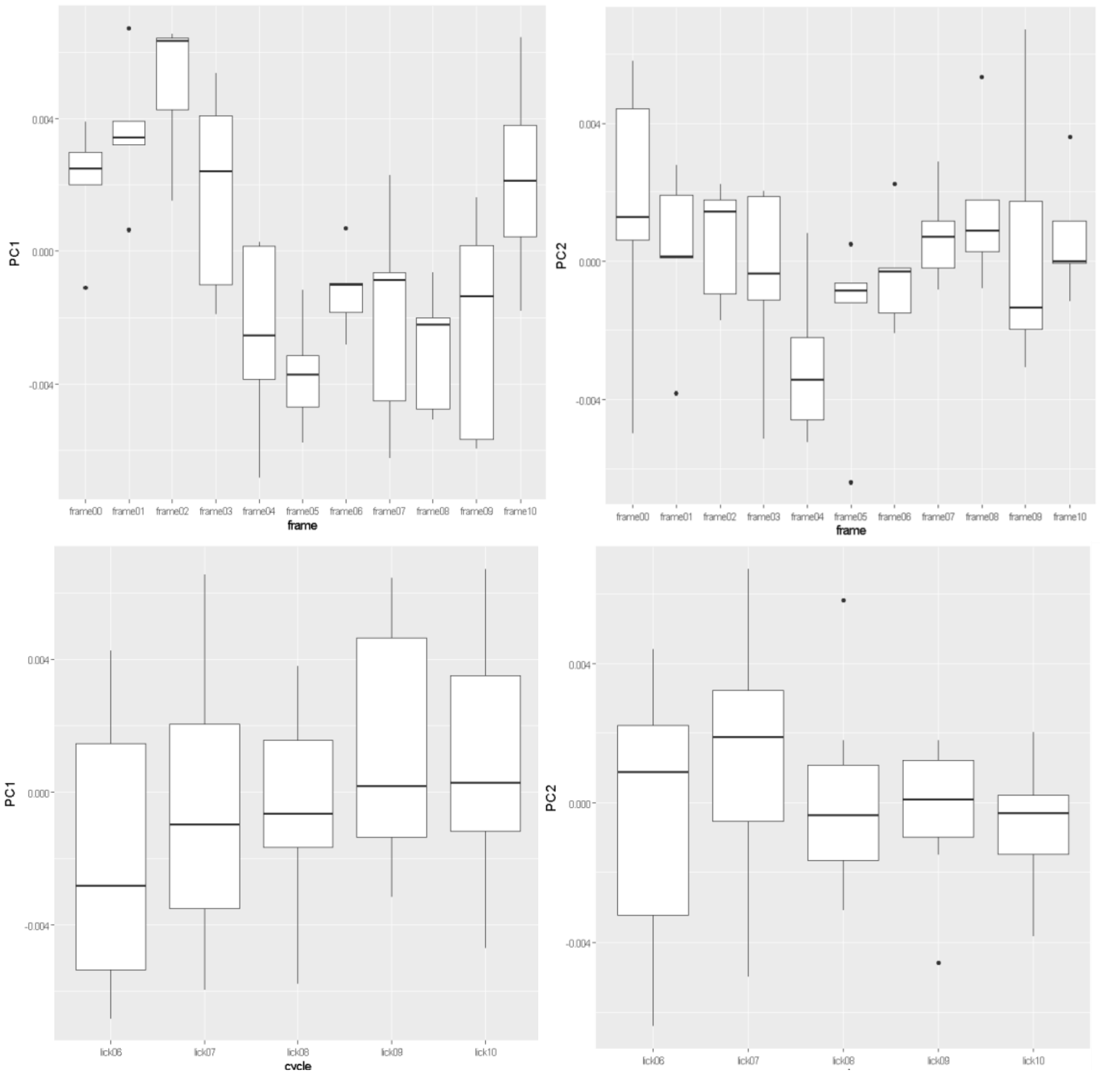


**Figure S6**. Primary sources of variation along the first two shape variables (PCs) that explain most of the variance in bill shape change for *Chalybura buffonii*. Series of boxplots showing variation among sequential bill landmark configurations, accounting for variation among lick cycles, along the PC axes (PC 1 left, PC 2 right) shown in figure 3 for *Chalybura buffonii*, by frame (pooled over five lick cycles; top row) and by lick cycles (pooled over frames, bottom row).

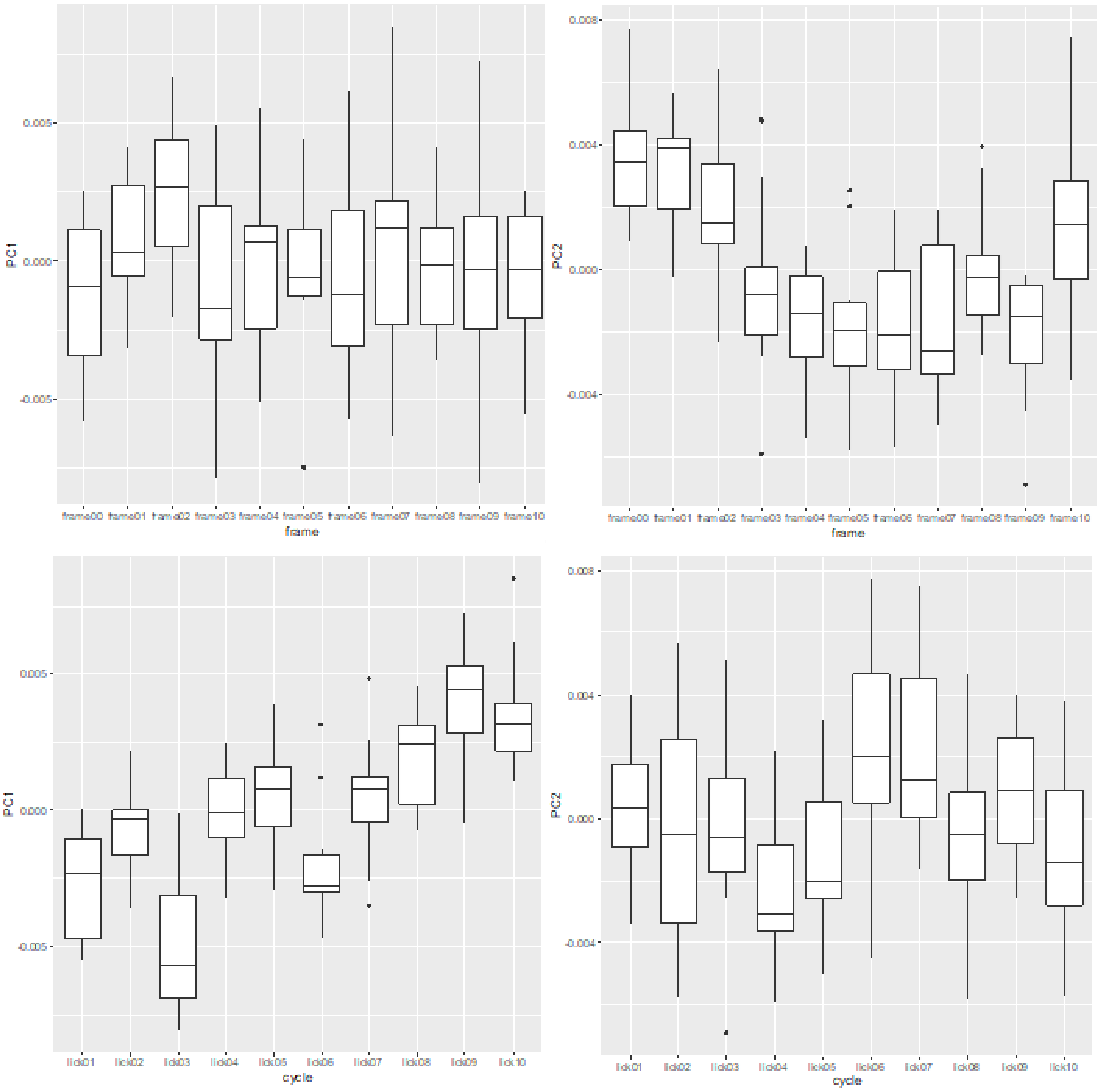


**Figure S7**. Primary sources of variation along the first two shape variables (PCs) that explain most of the variance in bill shape change for *Florisuga mellivora*. Series of boxplots showing variation among sequential bill landmark configurations, accounting for variation among lick cycles, along the PC axes (PC 1 left, PC 2 right) shown in figure 4 for *Florisuga mellivora*, by frame (pooled over five lick cycles; top row) and by lick cycles (pooled over frames, bottom row).

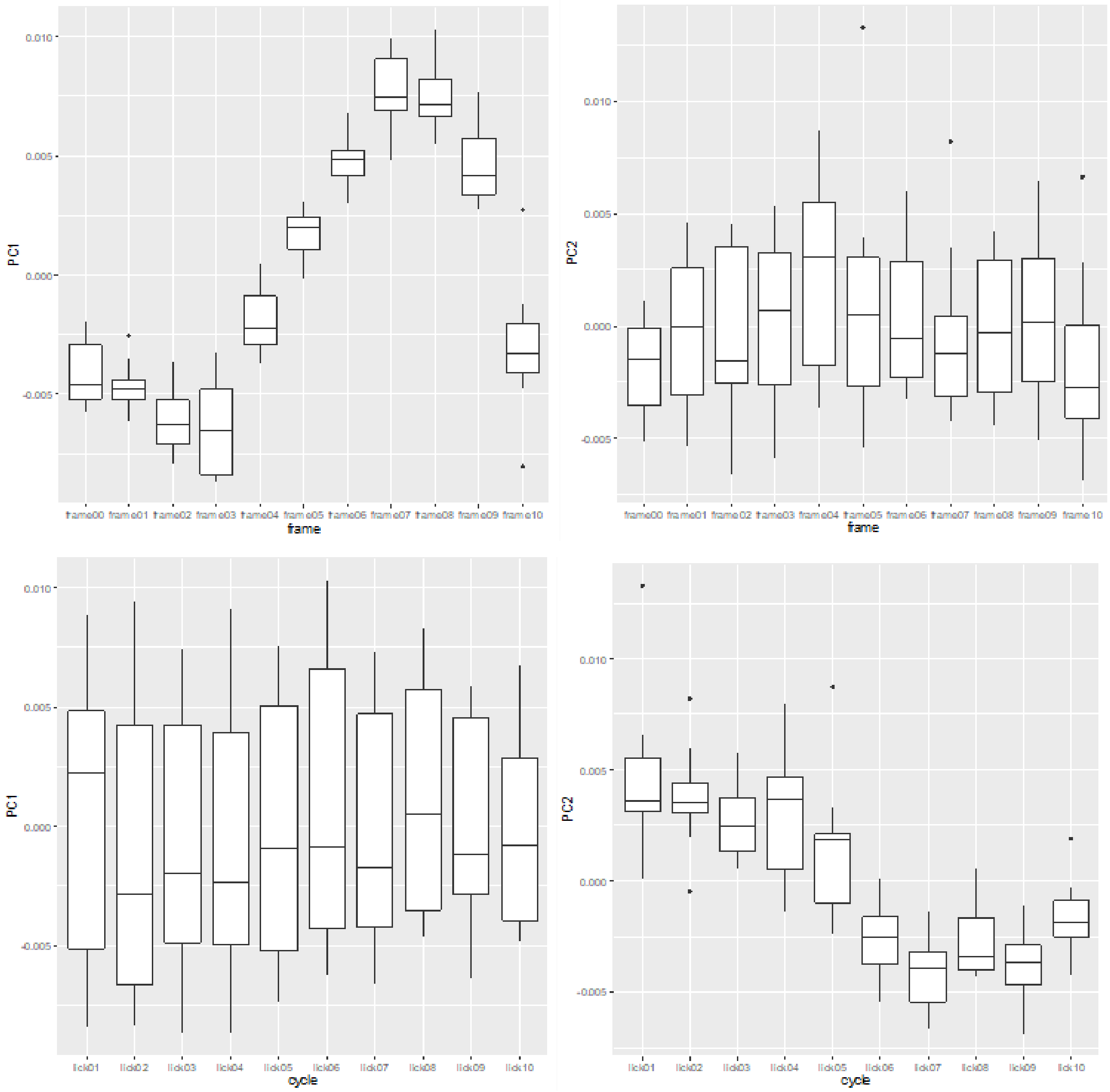


**Figure S8**. Primary sources of variation along the first two shape variables (PCs) that explain most of the variance in bill shape change for *Myrmia micrura*. Series of boxplots showing variation among sequential bill landmark configurations, accounting for variation among lick cycles, along the PC axes (PC 1 left, PC 2 right) shown in figure 4 for *Myrmia micrura*, by frame (pooled over five lick cycles; top row) and by lick cycles (pooled over frames, bottom row).

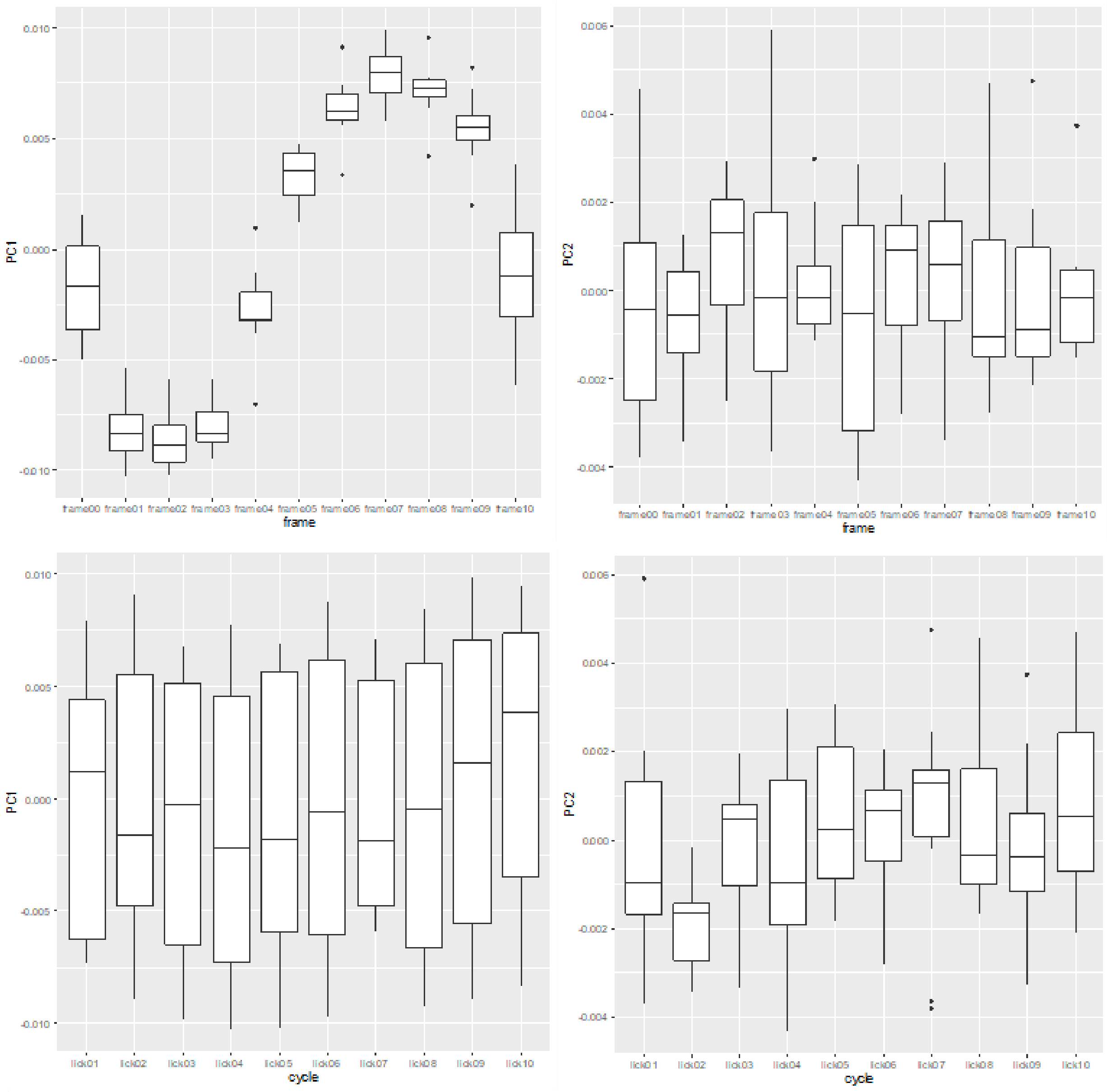


**Figure S9**. Primary sources of variation along the first two shape variables (PCs) that explain most of the variance in bill shape change for *Phaethornis longirostris*. Series of boxplots showing variation among sequential bill landmark configurations, accounting for variation among lick cycles, along the PC axes (PC 1 left, PC 2 right) shown in figure 4 for *Phaethornis longirostris*, by frame (pooled over five lick cycles; top row) and by lick cycles (pooled over frames, bottom row).

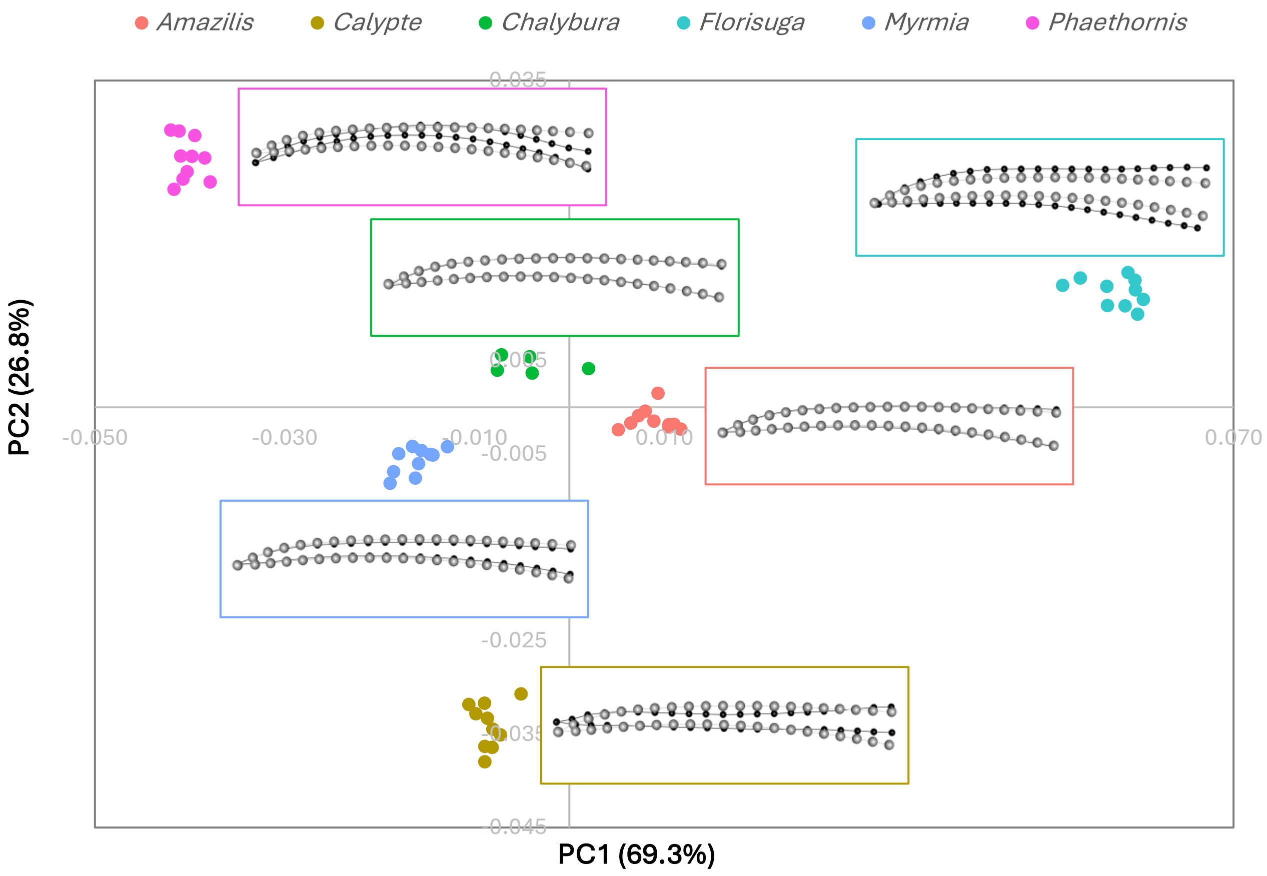


**Figure S10.** Overall (static) culmen shape variation among species in PC1-PC2 morphospace, based on the initial frames of each 5-10 lick cycles for each taxon. The insets show representative shape changes (black dots and lines; relative the grey dots and lines of the consensus configuration, magnified by 1.5X) from the centre of each species cluster of points.

**Figure S11**. Flexural rigidity profiles through the bills of (a) *Calypte anna*, and (b, c) two specimens of *Myrmia micrura* (cranial base of the bill left, rostral tip right). The light blue traces are the summed rigidities of the left and right *os maxillares*, and the pink trace depicts the rigidity of the *os nasale*. The dark blue trace shows the region of fusion between the latter two elements. Regions 1 and 3 (indicated by bold dashed vertical black lines) are areas of high flexural rigidity, whereas region 2 includes the "bending zone," characterized by low flexural rigidity, roughly corresponding to the intermediate part of the prepalatal upper jaw (see Fig. 5 in [8]).

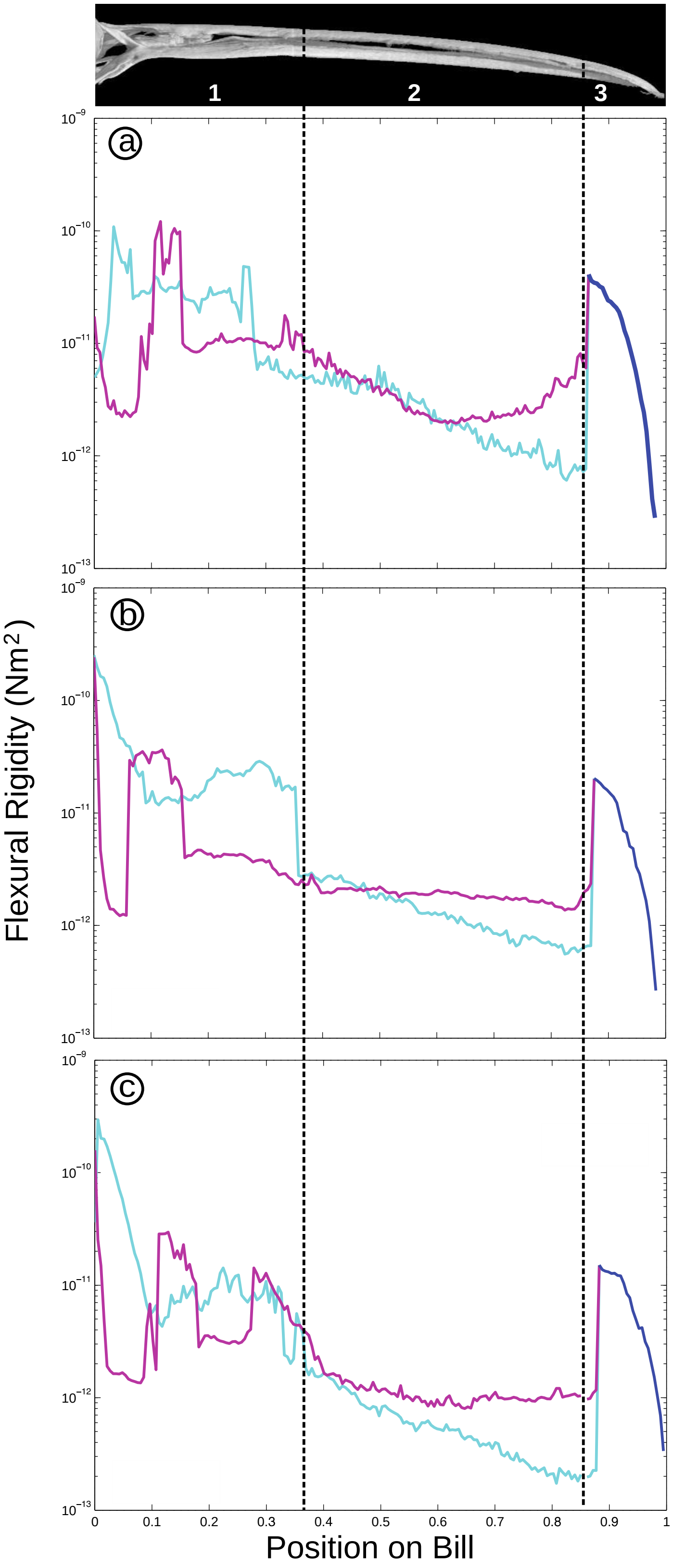

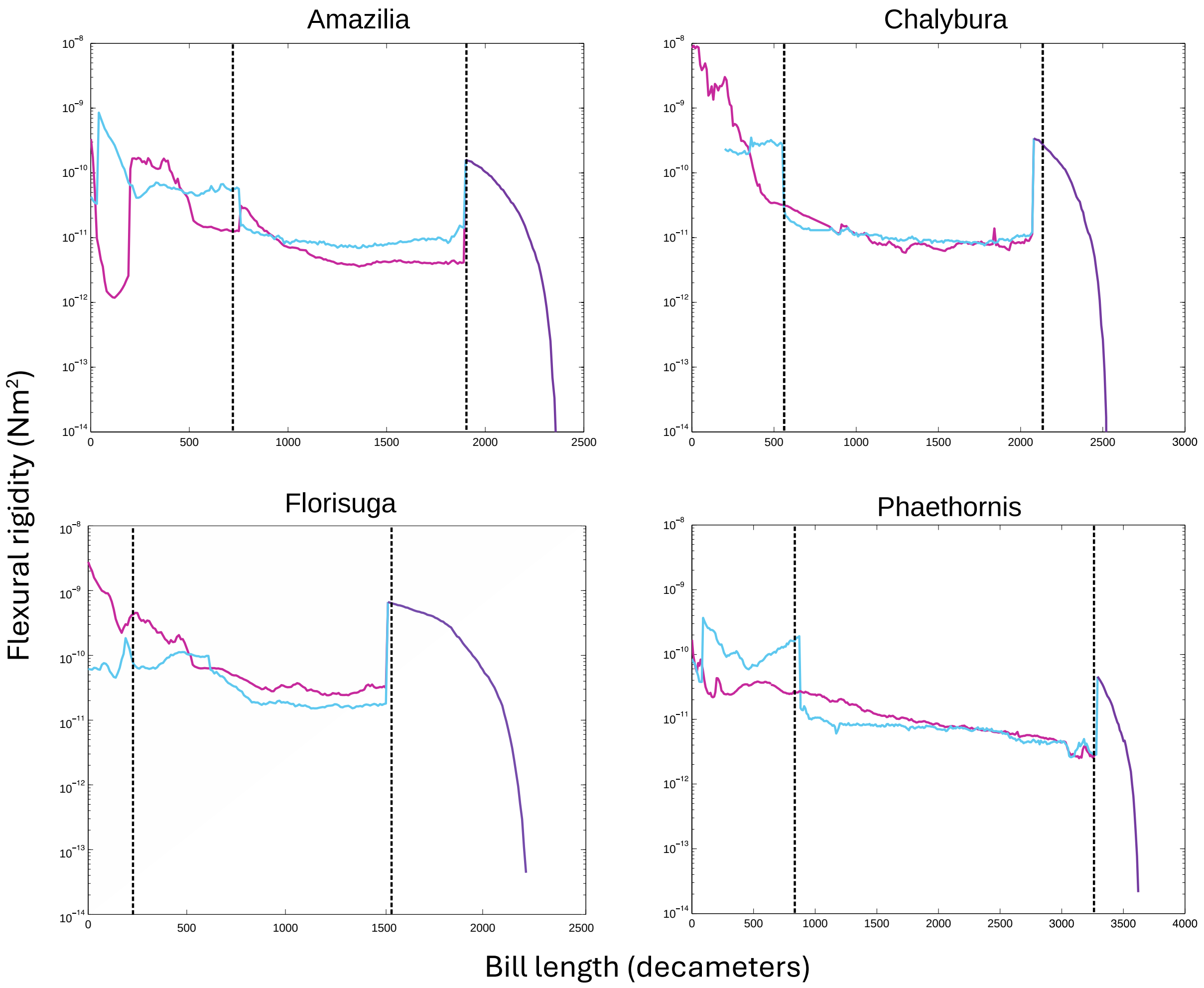


**Figure S12**. Flexural rigidity profiles through the bills of *Amazilis amazilia*, *Chalybura buffonii*, *Florisuga mellivora*, and *Phaethornis longirostris* (cranial base of the bill left, rostral tip right). The light blue traces are the summed rigidities of the left and right *os maxillares*, and the pink trace depicts the rigidity of the *os nasale*. The dark blue trace shows the region of fusion between the latter two elements. Regions 1 and 3 (indicated by bold dashed vertical black lines) are areas of high flexural rigidity, whereas region 2 includes the "bending zone," characterized by low flexural rigidity.

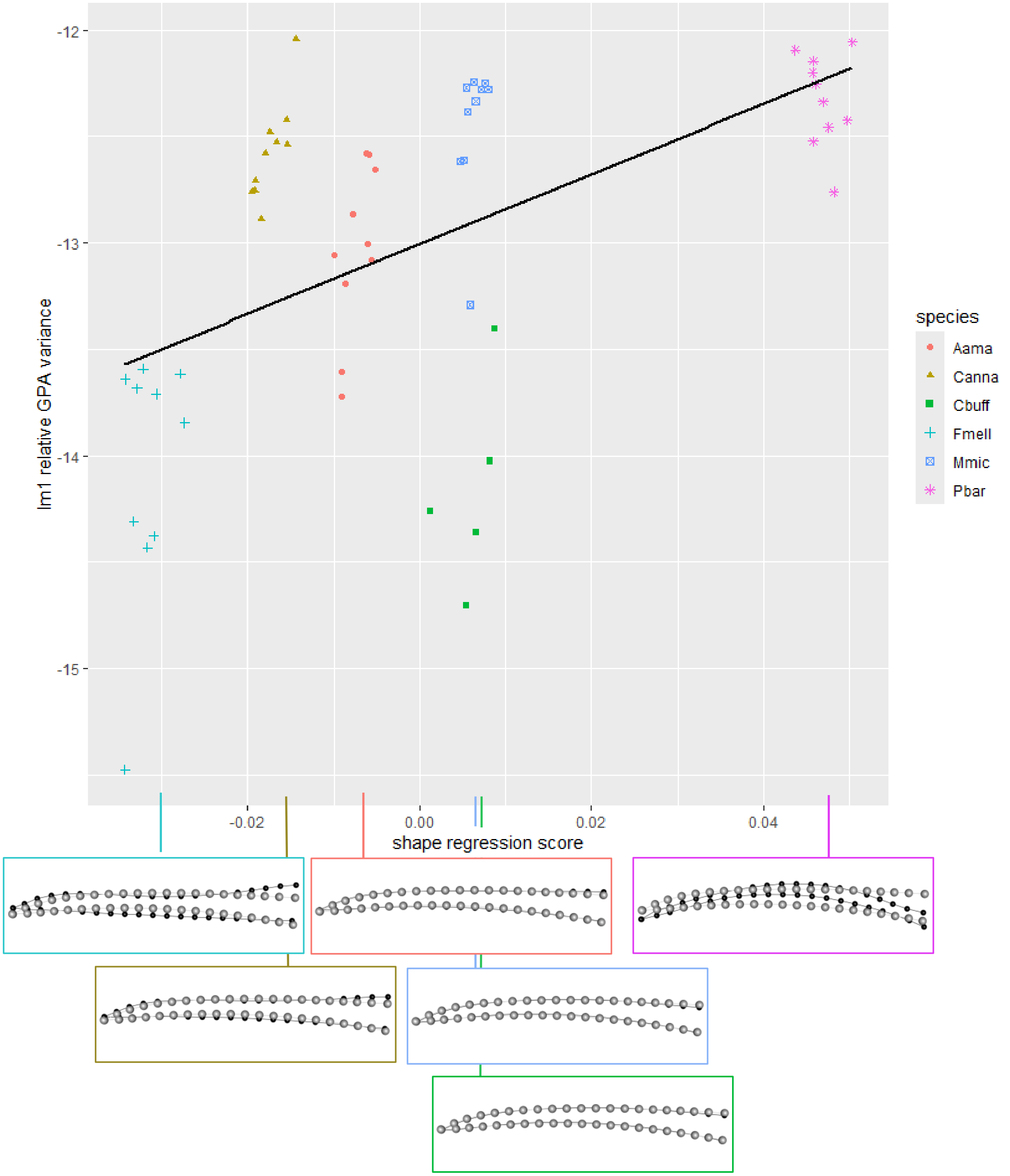


**Figure S13.** Scatterplot showing the relationship between the relative GPA variance of the distal bill tip landmark (LM 1) as a measure of the range of bill bending, and the shape regression scores, as a composite measure of bill shape and size (obtained from the covariation of shape and log centroid size [6,7]). The insets show shape differences (black dots and lines relative to the consensus configuration in grey, magnified by 3X) corresponding to the regression scores representative of each species along the x-axis.

**Table S1**. Results of a Procrustes ANOVA (R Package 'geomorph'; (68)) for *Calypte anna*, testing for bill shape differences among sequential frames of one lick cycle ("Lick02"), accounting for variation among repeated tracks ("rep") of the same cycle within each of five different participants ("part").

| | Df | SS | MS | Rsq | F | Z | Pr(>F) |
|---|---|---|---|---|---|---|---|
| part | 4 | 0.000987 | 0.00025 | 0.06922 | 9.0143 | 2.1884 | 0.0129 |
| frame | 10 | 0.009031 | 0.0009 | 0.63342 | 31.8806 | 11.1926 | 0.0001 |
| part:rep | 10 | 0.000274 | 2.74E-05 | 0.0192 | 0.9662 | -0.1083 | 0.544 |
| Residuals | 140 | 0.003966 | 2.83E-05 | 0.27816 | | | |
| Total | 164 | 0.014257 | | | | | |

Call: procD.lm(coords ~ part/rep + frame, iter = 9999, RRPP = TRUE, data = gdf); note that anova(fit, error = c("part:rep", "Residuals", "Residuals")) was used to appropriately test MS 'part' over MS 'part:rep', and MS 'part:rep' and MS 'frame' over residuals (from https://cran.r-project.org/web/packages/geomorph/vignettes/geomorph.assistance.html)

**Table S2**. Results of a Procrustes ANOVA (R Package 'geomorph'; [6]) for *Calypte anna*, testing for bill shape differences among sequential frames, accounting for variation among lick cycles.

| | Df | SS | MS | Rsq | F | Z | Pr(>F) |
|---|---|---|---|---|---|---|---|
| frame | 10 | 0.004682 | 0.00047 | 0.76791 | 45.9458 | 12.9758 | 0.0001 |
| cycle | 9 | 0.000498 | 5.53E-05 | 0.08167 | 5.4298 | 7.7801 | 0.0001 |
| Residuals | 90 | 0.000917 | 1.02E-05 | 0.15042 | | | |
| Total | 109 | 0.006097 | | | | | |

Call: procD.lm(coords ~ frame + dataset, iter=9999, data = gdf)

**Table S3**. Results of a Procrustes ANOVA (R Package ‘geomorph’; [6]) for *Amazilis amazilia*, testing for bill shape differences among sequential frames, accounting for variation among lick cycles.

| | Df | SS | MS | Rsq | F | Z | Pr(>F) |
|---|---|---|---|---|---|---|---|
| frame | 10 | 0.003626 | 0.00036 | 0.47439 | 10.4014 | 9.225 | 0.0001 |
| cycle | 9 | 0.00088 | 9.78E-05 | 0.11514 | 2.8051 | 6.5945 | 0.0001 |
| Residuals | 90 | 0.003138 | 3.49E-05 | 0.41047 | | | |
| Total | 109 | 0.007644 | | | | | |

Call: procD.lm(coords ~ frame + cycle, iter=9999, data = gdf)

**Table S4**. Results of a Procrustes ANOVA (R Package ‘geomorph’; [6]) for *Chalybura buffonii*, testing for bill shape differences among sequential frames, accounting for variation among lick five cycles.

| | Df | SS | MS | Rsq | F | Z | Pr(>F) |
|---|---|---|---|---|---|---|---|
| frame | 10 | 0.000672 | 6.72E-05 | 0.37403 | 2.8871 | 4.5573 | 0.0001 |
| cycle | 4 | 0.000194 | 4.84E-05 | 0.10777 | 2.0796 | 2.5231 | 0.0064 |
| Residuals | 40 | 0.000931 | 2.33E-05 | 0.51821 | | | |
| Total | 54 | 0.001796 | | | | | |

**Table S5**. Results of a Procrustes ANOVA (R Package ‘geomorph’; [6]) for *Florisuga mellivora*, testing for bill shape differences among sequential frames, accounting for variation among lick cycles.

| | Df | SS | MS | Rsq | F | Z | Pr(>F) |
|---|---|---|---|---|---|---|---|
| frame | 10 | 0.00105 | 1.05E-04 | 0.2004 | 3.3098 | 7.9236 | 0.0001 |
| cycle | 9 | 0.001334 | 1.48E-04 | 0.25465 | 4.6731 | 8.4177 | 0.0001 |
| Residuals | 90 | 0.002856 | 3.17E-05 | 0.54494 | | | |
| Total | 109 | 0.00524 | | | | | |

**Table S6**. Results of a Procrustes ANOVA (R Package 'geomorph'; [6]) for *Myrmia micrura*, testing for bill shape differences among sequential frames, accounting for variation among lick cycles.

| | Df | SS | MS | Rsq | F | Z | Pr(>F) |
|---|---|---|---|---|---|---|---|
| frame | 10 | 0.003214 | 0.00032 | 0.47326 | 13.4011 | 10.3828 | 0.0001 |
| cycle | 9 | 0.001419 | 0.00016 | 0.2089 | 6.5726 | 6.9328 | 0.0001 |
| Residuals | 90 | 0.002159 | 2.40E-05 | 0.31784 | | | |
| Total | 109 | 0.006792 | | | | | |

**Table S7**. Results of a Procrustes ANOVA (R Package 'geomorph'; [6]) for *Phaethornis longirostris*, testing for bill shape differences among sequential frames, accounting for variation among lick cycles.

| | Df | SS | MS | Rsq | F | Z | Pr(>F) |
|---|---|---|---|---|---|---|---|
| frame | 10 | 0.004236 | 0.00042 | 0.68096 | 22.6814 | 9.6993 | 0.0001 |
| cycle | 9 | 0.000304 | 3.38E-05 | 0.04884 | 1.8074 | 3.8898 | 0.0001 |
| Residuals | 90 | 0.001681 | 1.87E-05 | 0.27021 | | | |
| Total | 109 | 0.006221 | | | | | |

**Table S8**. Results of a Procrustes ANOVA (R Package 'geomorph'; [6]) testing for bill shape (i.e., the full set of Procrustes coordinates) and size (i.e., centroid size [Csize]) differences among species, based on the initial frames ("frame0") of 5-10 lick cycles for each taxon.

| | Df | SS | MS | Rsq | F | Z | Pr(>F) |
|---|---|---|---|---|---|---|---|
| species | 5 | 0.000123 | 2.47E-05 | 0.00159 | 0.9829 | -0.0068 | 0.5016 |
| Csize | 1 | 0.000062 | 6.18E-05 | 0.00079 | 2.4611 | 2.17926 | 0.0139 |
| species×Csize | 5 | 0.000155 | 3.10E-05 | 0.00199 | 1.2369 | 1.13527 | 0.13 |
| Residuals | 43 | 0.001079 | 2.51E-05 | 0.01387 | | | |
| Total | 54 | 0.077827 | | | | | |
| *Pooled interaction:* | | | | | | | |
| species | 5 | 0.051771 | 1.04E-02 | 0.6652 | 402.629 | 15.8565 | 0.0001 |
| Csize | 1 | 0.000194 | 1.94E-04 | 0.00249 | 7.5309 | 5.3132 | 0.0001 |
| Residuals | 48 | 0.001234 | 2.57E-05 | 0.01586 | | | |
| Total | 54 | 0.077827 | | | | | |

**Table S9**. Results of a general linear model testing the effects of bill shape and log centroid size encapsulated in shape regression scores (regscore; [6,7]) on the (log) GPA variance (relativized by the minimum variance observed during a given lick cycle) of the most rostral dorsal culmen landmark (LM 1), accounting for differences among species.

| Source | Df | SumSq | MeanSq | F-value | P-value |
|---|---|---|---|---|---|
| regscore | 1 | 9.8247 | 9.8247 | 77.8593 | < 0.0001 |
| species | 5 | 18.9837 | 3.7967 | 30.0886 | < 0.0001 |
| regscore x species | 5 | 0.6395 | 0.1279 | 1.0136 | 0.4215 |
| Residuals | 43 | 5.4260 | 0.1261 | | |
| Pooled interaction: | | | | | |
| regscore | 1 | 9.8247 | 9.8247 | 77.749 | < 0.0001 |
| species | 5 | 18.9837 | 3.7967 | 30.046 | < 0.0001 |
| Residuals | 48 | 6.0655 | 0.1264 | | |

Call: lm(formula = log(lm1relgpa) ~ regscore + species, data = df)

**Table S10. Hummingbird species included in the bill motion analyses.** We organise the species taxonomically to highlight the taxonomic breath of the sampling and order them according to bill length.

| Clade | Species | Bill length (mm) | Country |
|---|---|---|---|
| Topazes | *Florisuga mellivora* | 18.7 | Colombia |
| Hermits | *Phaethornis longirostris* | 40.2 | Ecuador |
| Emeralds | *Chalybura buffonii* | 23.3 | Colombia |
| | *Amazilis amazilia* | 19.4 | Colombia |
| Bees | *Calypte anna* | 16.3 | USA |
| | *Myrmia micrura* | 15.7 | Ecuador |
